# Persistent virus-specific and clonally expanded antibody secreting cells respond to induced self antigen in the CNS

**DOI:** 10.1101/2022.08.29.505678

**Authors:** Andreas Agrafiotis, Raphael Dizerens, Ilena Vincenti, Ingrid Wagner, Raphael Kuhn, Danielle Shlesinger, Marcos Manero-Carranza, Tudor-Stefan Cotet, Kai-Lin Hong, Nicolas Page, Nicolas Fonta, Ghazal Shammas, Alexandre Mariotte, Margot Piccinno, Mario Kreutzfeldt, Benedikt Gruntz, Roy Ehling, Alessandro Genovese, Alessandro Pedrioli, Andreas Dounas, Sören Franzenburg, Vladyslav Kavaka, Lisa Ann Gerdes, Klaus Dornmair, Eduardo Beltrán, Annette Oxenius, Sai T. Reddy, Doron Merkler, Alexander Yermanos

## Abstract

B cells contribute to the pathogenesis of both cellular- and humoral-mediated central nervous system (CNS) inflammatory diseases through a variety of mechanisms. In such conditions, B cells may enter the CNS parenchyma and contribute to local tissue destruction. It remains unexplored, however, how infection and autoimmunity drive transcriptional phenotypes, repertoire features, and antibody functionality. Here, we profiled B cells from the CNS of murine models of intracranial (i.c.) viral infections and autoimmunity. We identified a population of clonally expanded, antibody secreting cells (ASCs) that had undergone class-switch recombination and extensive somatic hypermutation following i.c. infection with attenuated lymphocytic choriomeningitis virus (rLCMV). Recombinant expression and characterisation of these antibodies revealed specificity to viral antigens (LCMV glycoprotein GP), correlating with ASC persistence in the brain weeks after resolved infection. Furthermore, these virus-specific ASCs upregulated proliferation and expansion programs in response to the conditional and transient induction of the LCMV GP as a neo-self antigen by astrocytes. This class-switched, clonally expanded, and mutated population persisted and was even more pronounced when peripheral B cells were depleted prior to autoantigen induction in the CNS. In contrast, the most expanded B cell clones in mice with persistent expression of LCMV GP in the CNS did not exhibit neo-self antigen specificity, potentially a consequence of local tolerance induction. Finally, a comparable population of clonally expanded, class-switched, proliferating ASCs was detected in the cerebrospinal fluid of multiple sclerosis patients. Taken together, our findings support the existence of B cells that populate the CNS and are capable of responding to locally encountered autoantigens.

**Graphical abstract:** 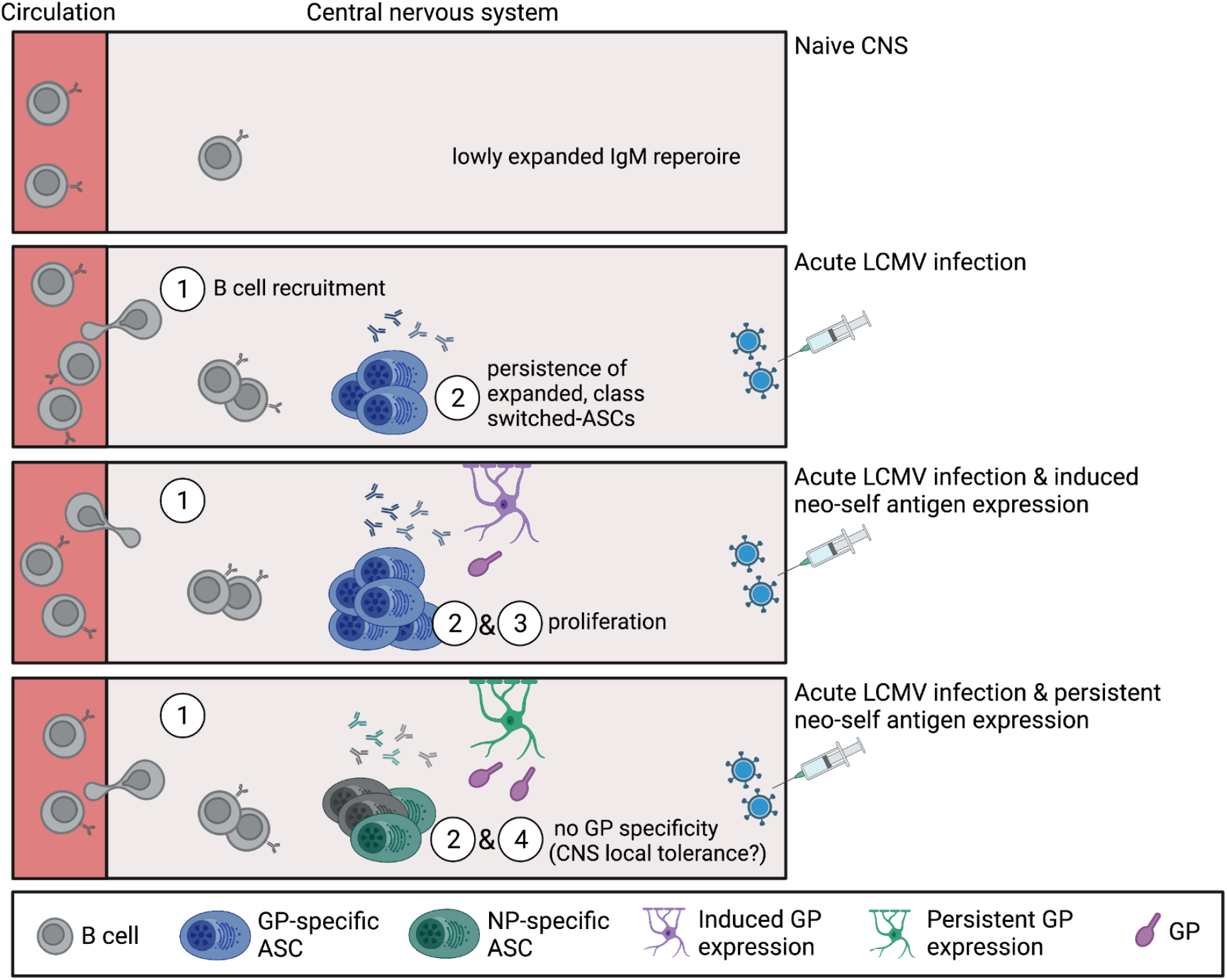

## Introduction

The access of circulating immune cells into the CNS is restricted in healthy individuals under steady-state conditions with only very low numbers of lymphocytes residing in the brain. However, in CNS inflammatory conditions, the number of immune cells entering the CNS parenchyma through the blood-brain barrier or blood-cerebrospinal fluid barrier, increases dramatically (Wilson, Weninger, and Hunter 2010; Schwartz et al. 2013; Prinz and Priller 2017; Agrawal et al. 2006; Owens, Bechmann, and Engelhardt 2008; Ghersi-Egea et al. 2018). Upon infiltration, T and B cells can adopt a wide range of phenotypes and effector functions that can either protect against pathogenic threat or, conversely, contribute to disease (Frebel, Richter, and Oxenius 2010; Li, Patterson, and Bar-Or 2018; Jain and Yong 2021). Thereby, T cells have previously been shown to be able to locally reside in a variety of non-lymphoid tissues, including the CNS, and to adopt a tissue-resident memory phenotype (Trm)(Jiang et al. 2012; Schenkel and Masopust 2014; Carbone 2015; Clark 2015; Dijkgraaf et al. 2019). Such Trm are poised to respond to secondary exposures (Steinbach et al. 2016; Urban et al. 2020; Vincenti et al. 2022) or can participate in compartmentalized inflammation in the CNS (Vincenti et al. 2022), In contrast, much less is known about the phenotype and function of tissue-resident B cells (Allie and Randall 2020; Gray and Farber 2022) and their interference with circulating counterparts. This is particularly true in the case of the CNS, where the formation and existence of resident B cells is still under debate. Yet, B cells are involved in the pathogenesis of neuroinflammatory diseases by a variety of mechanisms including antigen presentation to T cells (Meinl, Krumbholz, and Hohlfeld 2006), transport of antigens to secondary lymphoid organs, secretion of pro-inflammatory or anti-inflammatory cytokines (Fillatreau et al. 2002; Barr et al. 2012), and pathogenic antibodies (Sabatino, Pröbstel, and Zamvil 2019; Jain and Yong 2021). Furthermore, the success of B cell-depleting therapies in patients with multiple sclerosis and other chronic autoimmune conditions with presence of neuronal auto-antibodies provides compelling evidence that B cells are crucially involved in the pathophysiology of these diseases (Krumbholz et al. 2012; Chitnis and Weiner 2017; Lee, Rojas, and Gommerman 2021; Häusser-Kinzel and Weber 2019; Smets and Titulaer 2022). While autoreactive antibodies to various protein targets such as MOG (Brilot et al. 2009), DPPX (Boronat et al. 2013), and aquaporin-4 (Jarius et al. 2008) have been profiled and are even used in diagnostics of certain neurological disorders (Dalakas, Alexopoulos, and Spaeth 2020; Prüss 2021), the cellular counterparts and origins within or outside the CNS of these antibodies remain unclear. In addition, despite recent advances in the field regarding transcriptional characteristics of disease-relevant B cells subsets found in the inflamed CNS (Ramesh et al. 2020), whether these cells can undergo clonal selection and be locally reactivated to CNS-restricted autoantigens remains unknown.

Recent advancements in sequencing and microfluidic techniques have enabled comprehensive profiling of B cells and their corresponding antibody repertoires (Yermanos, Agrafiotis, et al. 2021; Neumeier et al. 2021; Agrafiotis et al. 2021; Horns, Dekker, and Quake 2020). Current iterations of this technology provide both single-cell antibody repertoires (full-length, paired heavy and light chain antibody sequences) and single-cell transcriptomes (Csepregi et al. 2020). This results in a quantitative profile of B cell clonal selection that can further be validated via recombinant expression of antibodies of interest. However, the relationship between gene expression and B cell receptor (BCR) repertoire of CNS B cells has not yet been linked to the functional properties of their corresponding antibodies. Here, we leveraged single-cell antibody repertoire and transcriptome sequencing in combination with transgenic murine models of infection and autoimmunity to investigate the clonal selection of virus-specific B cells. We discovered populations of antigen-specific ASCs that had undergone class-switch recombination, clonal expansion, and somatic hypermutation after viral infection in the CNS. Upon induction of localized LCMV GP as a neo-self antigen in glia cells after viral clearance, we observed that ASCs adopted a proliferative transcriptional program and produced antibodies specific to the induced neo-self antigen. In contrast, such GP-specific antibodies were absent after a corresponding transient viral infection in which LCMV GP was persistently expressed as a neo-self antigen in oligodendrocytes,, pointing towards a CNS-mediated tolerance mechanism. Mirroring our experimental observations, we uncovered expanded and class-switched ASCs in the cerebrospinal fluid of patients with relapsing multiple sclerosis suggesting a disease-relevant involvement of ASC in CNS autoimmunity which may serve as a starting point for future specific therapeutic approaches.

## Results

### Models of inducible and chronic expression of CNS-localized neo-self antigen

To characterize virus-specific and potentially autoreactive B cells in the CNS, we utilized four different experimental conditions involving viral infection and autoimmunity in the CNS (Figure 1A). Three conditions utilized a recently described CNS autoimmune mouse model driven by brain-resident memory T cells (bTRM) (Vincenti et al. 2022), which involves GFAP-Cre^ERT2 tg/+^;Stop-GP^flox/+^ mice (referred to as GFAP:GP). GFAP:GP mice express a tamoxifen-inducible Cre recombinase under the glial fibrillary acidic protein (GFAP) promoter driving the expression of LCMV glycoprotein (GP). The GFAP:GP model generates LCMV-GP33-41-specific bTRM by an intracranial infection with rLCMV-GP58, a recombinant LCMV strain containing the first 58 amino acids of LCMV GP fused to the vesicular stomatitis virus (VSV) glycoprotein (termed rLCMV-GP58) (Figure 1A). Six weeks after viral clearance, intraperitoneal (i.p.) tamoxifen (TAM) administration induces full-length LCMV-GP as a neo-self antigen in astrocytes and results in bTRM reactivation and development of locomotor deficits (Vincenti et al. 2022). Littermate Stop-GP^flox/+^ mice, termed STOP:GP, served as post-infection controls as they lack the GFAP-Cre^ERT2tg/+^ and therefore do not express the full-length LCMV GP upon TAM administration (Vincenti et al. 2022) (Figure 1A). To exclude recruitment of peripheral B cells during induced neo-self antigen expression (LCMV-GP), we included an experimental condition that received i.p. administration of a B cell CD20-depleting monoclonal antibody before administration of TAM (Figure S1A), referred to as aCD20-GFAP:GP (Figure 1A). Lastly, we leveraged recently described transgenic mice that constitutively express the full-length LCMV GP under the control of the myelin oligodendrocyte glycoprotein (MOG) promoter, termed MOG:GP, to create a condition in which viral infection induces chronic autoimmune disease (Page et al. 2021) (Figure 1A). Mice of the GFAP:GP, aCD20-GFAP:GP, and STOP:GP conditions were sacrificed seven days after initial TAM administration, while mice of MOG:GP were sacrificed 47 days after initial infection. Mice started to lose weight approximately three days following neo-self antigen induction in all groups excluding STOP:GP mice (Figure 1B). Furthermore, all groups losing weight demonstrated locomotor impairments and ataxia based on reduced performance on the rotarod test (Figures 1B). Together, this revealed that circulating CD20-expressing B cells are not required for the observed disease in the previously described GFAP:GP model of autoimmunity despite histology revealing a significant increase in B220+ cell number in brains and spinal cords of GFAP:GP mice compared to naive mice (Figures 1C, 1D).

**Figure 1.**
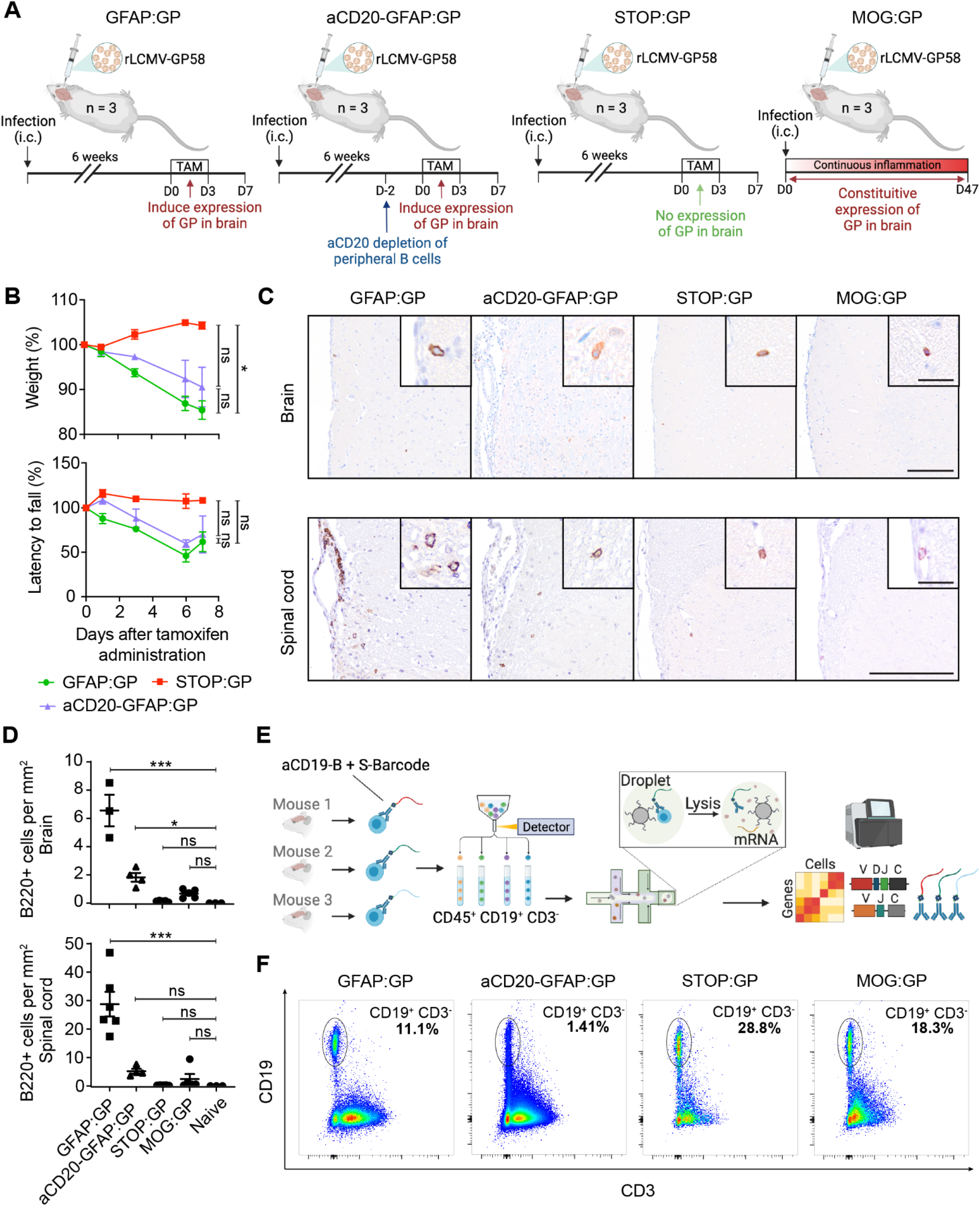
Models of inducible and chronic expression of CNS-localized self-antigen. A. Experimental overview of intracranial (i.c.) infection and models of autoimmunity. B. Weight loss (top) and rotarod performance measuring locomotor impairment and ataxia (bottom), where data represent means ± SEM. *P < 0.05; ns, not significant; two-way ANOVA followed by Tukey’s multiple comparisons test. C. Representative immunostainings and D. quantification of B220+ cells in brain and spinal cord sections of indicated groups. Tissues were collected 7 days after tamoxifen treatment, except for MOG:GP group, where tissue collection occurred 50 days post infection. Scale: 150um. Inset: 20um. Symbols represent one individual mouse, and bars represent means ± SEM. *P < 0.05, ***P < 0.001; ns, not significant; one-way ANOVA followed by Dunnett’s multiple comparisons test. E. Fluorescent activated cell sorting to isolate CD19^+^ expressing cells from murine brains followed by library construction and single-cell sequencing. F. Flow cytometry dot plot analysis of sorted CD19^+^/CD3^-^ B cell populations per mouse for each experimental group. One representative mouse per group is shown with the percentage of the sorted population indicated.

### Single-cell immune repertoire sequencing reveals heterogeneous populations of CNS B cells following infection and autoimmunity

Given previous findings demonstrating B cell infiltration in GFAP:GP, STOP:GP and MOG:GP conditions (Vincenti et al. 2022; Page et al. 2021), in conjunction with our observed disease manifestations (Figures 1B), we questioned whether single-cell immune repertoire sequencing would provide insight into selection fingerprints of CNS B cells and their functional role in the context of CNS infection and autoimmunity. We therefore isolated brain-infiltrating B cells using flow cytometry using anti-CD19 antibodies tagged with mouse-specific oligonucleotide barcodes (Figure 1E). Sorted CD19^+^/CD3^-^ B cells (Figures 1F, S1B) from three mice per group were pooled for each of the aforementioned experimental conditions (Figure 1A) and subjected to single-cell sequencing of their antibody repertoires, transcriptomes, and mouse-specific oligonucleotide barcodes (Figure 1E). Following library construction, deep sequencing, and alignment to reference genomes, we recovered a total of 22,444 single cells per condition with an average of 766 genes per cell and a low percentage (~1-3%) of reads mapping to mitochondrial genes (Figures S2A, S2B, S2C). The fractions of cells assigned to each mouse-specific oligonucleotide barcode were comparable within all experimental conditions for both the transcriptomes and antibody repertoires with only a minor fraction (~5%) of cells that could not be assigned to any of the barcodes (Figure S2D), thereby signifying high reproducibility across mice.

We performed uniform manifold approximation projection (UMAP) and unsupervised clustering for an unbiased overview of all cell populations based on global gene expression (Figure 2A, S3). This revealed that cells arising from STOP:GP and MOG:GP mice occupied multiple clusters at comparable frequencies, indicating that brain-infiltrating B cells after long-term CNS autoimmunity are transcriptionally similar to steady-state B cells generated via infection. Cells from GFAP:GP were present in the majority of observed clusters but were especially present in cluster 0 (Figures 2B, 2C), whereas cells arising from the aCD20-GFAP:GP group were largely restricted to clusters 2, 5, and 9 (Figures 2B, 2C). Together this highlights the effects of peripheral B cell depletion before the induction of LCMV GP expression in astrocytes and suggests that cluster 0 is most derived from peripherally recruited cells whereas clusters 2, 5, and 9 are not recirculating persisting in the CNS.

**Figure 2.**
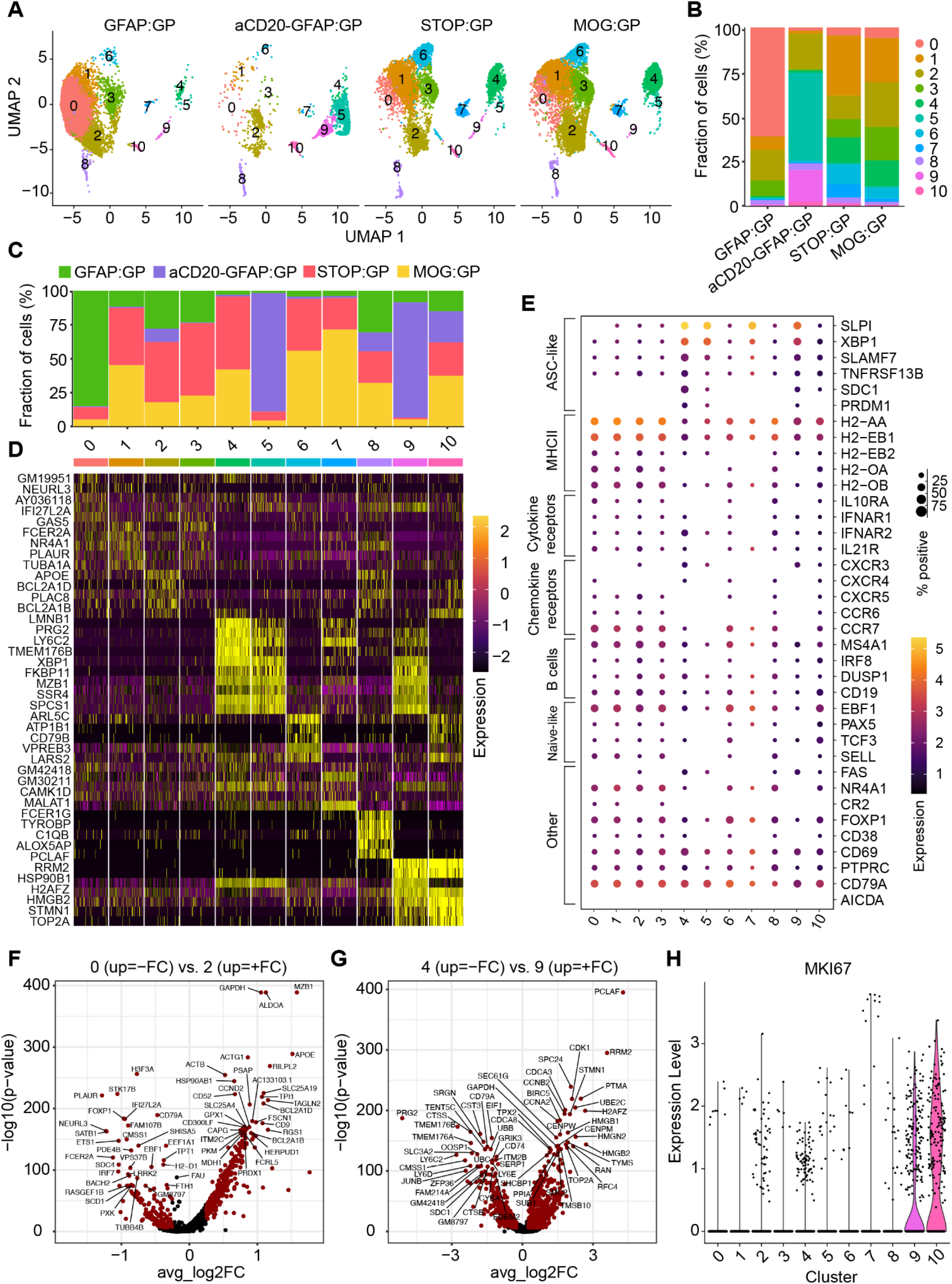
Single-cell immune repertoire sequencing reveals heterogeneous populations of CNS B cells following infection and autoimmunity. A. Uniform manifold approximate projection (UMAP) split by sample and colored by transcriptional cluster. B. The fraction of cells in each transcriptional cluster separated by experimental group. C. Distribution of experimental groups per transcriptional cluster. D. Differentially expressed genes defining each cluster. E. Gene expression levels of B cell markers separated by cluster membership (x axis). The size of the dot corresponds to the percentage of cells expressing the given marker and the color indicates the mean expression per cell within each cluster. F. Differentially expressed genes between clusters 0 and 2. Points in red indicate differentially expressed genes (adjusted p-value < 0.01 and average log2 fold change (FC) > 0.25) G. Differentially expressed genes between clusters 4 and 9. H. Normalized expression of *MKI67* separated by transcriptional cluster.

### CNS infection generates long-lived brain-infiltrating ASCs that persist during autoimmunity

To further characterize B cell phenotypic diversity across all experimental conditions, we performed differential gene analysis, gene ontology, and gene set enrichment for each of the transcriptional clusters. Cluster 0 expressed genes related to interferon and antigen-presentation and was almost entirely composed of cells arising from GFAP:GP mice following TAM-mediated induction of GP and almost entirely absent in aCD20-GFAP:GP mice (Figures 2C, 2D, S4A, S5). Cells from cluster 2 expressed genes related to germinal center B cells and contained comparable fractions of B cells from all four groups (Figures 2C, 2D, 2E, S4B), representing a cluster that was likely induced by rLCMV-GP58 infection and was independent of GP neo-self antigen expression. Cells from clusters 4, 5 and 9 expressed genes related to ASCs (Figure 2D, S4B, S5). Visualizing the expression of known B cell markers confirmed the unbiased transcriptional properties that ASC-phenotype of clusters 4, 5, and 9, as they expressed genes associated with antibody secretion programs (e.g., *Cd138* (*Sdc1*), *Taci* (*Tnfrsf13b*), *Xbp1, Slamf7, Nur77*) (Figures 2E, S6). Cells from GFAP:GP mice expressed higher levels of genes associated with naive B cells and antigen-presentation (*Cd19, Sell, Ccr7*, and MHCII genes) (Figures 2E, S6). The observed ASC and GC B cell phenotypes were in stark contrast to the transcriptional profiles of previously profiled CD19+ B cells populating the CNS in naive and aged mice which entirely lacked ASC-signatures (Yermanos, Neumeier, et al. 2021). Together, this implies that i.c. infection with rLCMV-GP58 induced ASCs in the brain that persist weeks following viral clearance (STOP:GP group) and that are maintained during autoimmunity (aCD20-GFAP:GP and MOG:GP groups).

As cluster 2 did not correspond to an ASC-phenotype yet remained after peripheral B cell depletion (~25% of all B cells from aCD20-GFAP:GP located in cluster 2 (Figure 2B)), we therefore calculated the differentially expressed genes between clusters 0 and 2 to shed light on those cells highly-specific to the GFAP:GP group. This demonstrated that genes such as *Apoe, Mzb1, Ccnd2*, and genes related to *Bcl2*, were upregulated in cluster 2, whereas genes such as MHCII genes were relatively upregulated in cluster 0 (Figures 2F, S4A). Together with the detected expression of *Fad* and *Cxcr3*, this suggests that TAM-driven induction of astrocytic GP recruits peripheral B cells that contribute in antigen-presentation and that B cells with upregulated germinal center markers persist in the CNS following peripheral B cell depletion.

### Peripheral B cell depletion enriches proliferating ASCs upon neo-self antigen induction

Having observed that ASC phenotypes were present across multiple clusters (4, 5, and 9) in a group-specific manner (Figure 2A), we next questioned the underlying transcriptional heterogeneity of these subsets. While MOG:GP and STOP:GP mice contained cells located predominantly in cluster 4, the B cells isolated from aCD20-GFAP:GP mice were preferentially located in clusters 5 and 9 (Figure 2B), suggesting that the inducible expression of LCMV GP by astrocytes further differentiated these cells. Of note, this ASC-phenotype is also visible in cells from GFAP:GP group, although likely masked due to massive recruitment of peripheral B cells that do not belong to clusters 4, 5 and 9. Expression of CD20 could still be detected within brain-derived cells from the aCD20-GFAP:GP group, albeit at a lower frequency (Figure S7), suggesting that these cells were located behind the blood brain barrier and thus inaccessible for depleting antibodies in the circulation. We next hypothesized that the B cells could have potentially adopted distinct transcriptional phenotypes due to the inflammatory milieu present following the induction and expression of GP by astrocytes. Performing differential gene expression analysis between clusters 4 and 9 revealed that cell-cycle and proliferation genes were enriched in cluster 9, which paralleled the expression of the proliferation marker KI67 (MKI67) (Figure 2H). Similarly, we observed cell proliferation associated genes, such as *Sub1* and *Fkbp11* (Chakravarthi et al. 2016; Qiu et al. 2021), enriched in cluster 5 relative to cluster 4 and less pronounced in cluster 9 (Figure S8A, S8B), which further suggests a proliferative phenotype of the ASCs arising from aCD20-GFAP:GP brains.

### Brain ASCs are clonally expanded and class-switched following viral infection and induction of autoimmunity

After having observed ASC expression profiles of B cells following infection and autoimmunity, we next questioned whether integrating antibody repertoire sequencing information would help elucidate the selection histories experienced by these B cells. We could recover full-length, paired-heavy-light chain antibody sequences for approximately 12,000 B cells, which resulted in a total of 8,187 clones across all experimental groups. Visualizing the fraction of clones that were supported by two or more unique cell barcodes demonstrated that clonal expansion was only detected in 17% of cells in the GFAP:GP group (Figure 3A), supporting a model where peripheral B cells infiltrate in an antigen-independent manner. This was in stark contrast to the condition in which peripheral B cells were depleted, as clonal expansion was detected in 77% of clones (Figure 3A). This high degree of clonal expansion in aCD20-GFAP:GP mice relative to other groups hinted towards a local antigen-reencounter event and local proliferation during the induction of GP following TAM administration, as baseline clonal expansion was detected in 45% of clones in STOP:GP mice (Figure 3A).

**Figure 3.**
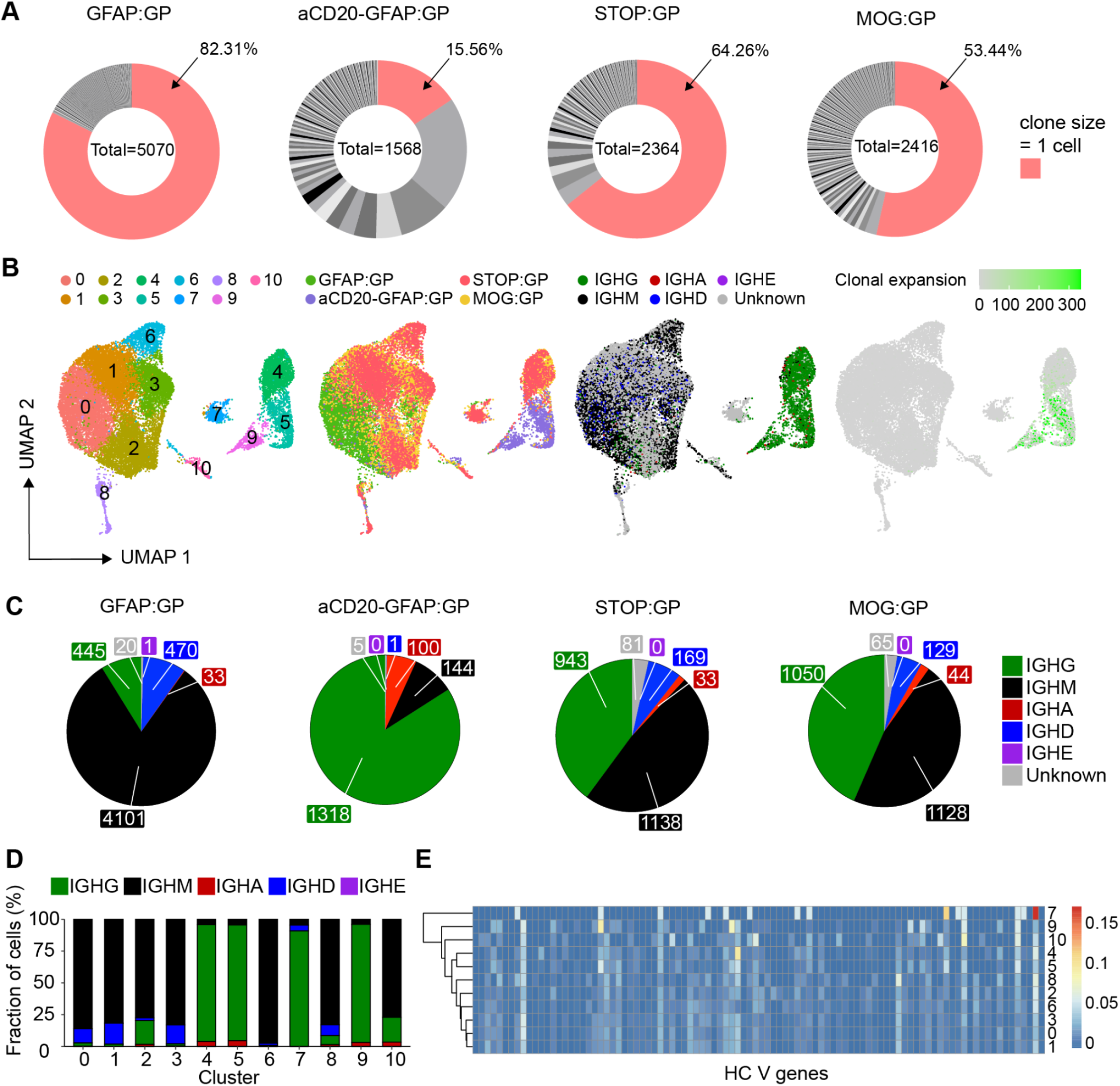
Profiling the clonally expanded, class-switched antibody secreting cells (ASCs) populating the brain following intracranial infection. A. Distribution of clonal expansion for each experimental group. Each section corresponds to a unique clone and the size corresponds to the fraction of cells relative to the total repertoire. Red color highlights the fraction of clones containing 1 cell. The number in the center of the circle refers to the total number of recovered B cells. B. Uniform manifold approximation projection (UMAP) depicting transcriptional cluster, group identity, antibody isotype, and clonal expansion for all recovered single-cells. Clonal expansion refers to the number of cells assigned to a single clone based on default clonotyping using Enclone. C. Distribution of antibody isotype for all cells within each experimental group. D. Distribution of antibody isotype for all cells within each transcriptional cluster filtering Cells with no isotype assigned (unknown) were removed. E. Heavy chain clonal V gene usage across transcriptional clusters.

To provide a more detailed description of these clonally expanded B cells, we next integrated repertoire features with transcriptional information. When overlaying antibody isotype onto the previously computed UMAP (Figure 2A), we discovered that the ASCs like B cells in clusters 4, 5, and 9 were IgG-expressing and had higher levels of clonal expansion relative to the remaining IgM-dominant clusters (Figure 3B). When quantifying isotype usage, we observed that GFAP:GP mice had the least amount of IgG, whereas this isotype represented the highest proportion for aCD20-GFAP:GP mice (Figure 3C, 3D). In contrast, a considerable fraction (~50%) of B cells expressed IgG in STOP:GP and MOG:GP mice (Figure 3C, 3D). The presence of these cells in MOG:GP mice shows that a population of clonally expanded, IgG-expressing ASCs cells persist in the brain 7 weeks following i.c. viral infection despite persistent neo-self antigen expression. Having observed that certain B cell phenotypes were dominated by IgG-producing cells, we questioned whether these clusters preferentially employed certain germline genes, as this could hint towards shared antigen exposure. Indeed, hierarchical clustering demonstrated that ASC clusters (4, 5, 9) diverged from those clusters lacking the ASC phenotype (Figure 3E), suggesting common selective pressures for those clonally expanded and IgG-expressing ASCs. Interestingly, this is in contrast to prior studies that have demonstrated IgA-secreting plasma cells in the meninges during homeostasis (Fitzpatrick et al. 2020).

### Clonally expanded and class-switched ASCs in the CNS are virus-specific and modulated by neo-self antigen induction

After characterizing the repertoire and transcriptional properties of clonally expanded ASCs cells, we investigated the functional properties of their antibodies. Visualizing the most expanded clones for each group demonstrated that despite GFAP:GP, STOP:GP, and MOG:GP groups having significant fractions of IgM-expressing cells (Figure 3C, 3D), the vast majority of the most expanded clones were exclusively of the IgG isotype (Figure S9A). Furthermore, we detected considerable somatic hypermutation within these expanded clones, as demonstrated by clonally related B cells expressing different antibodies (Figure 4A). To discover the antigenic targets of these expanded IgG-producing B cells, we recombinantly expressed the antibodies from the most expanded clones for each group and validated antigen-specificity using enzyme-linked immunosorbent assay (ELISA) against a panel of LCMV, VSV, and unrelated antigens. We discovered that multiple clones arising from GFAP:GP, aCD20-GFAP:GP, and STOP:GP were specific to LCMV GP, whereas only LCMV nucleoprotein (NP) and no LCMV GP specificity was detected in the MOG:GP group (Figure 4B, S9B, S10). While the GP- and NP-binding clones demonstrated distinct evolutionary histories, CDR3 sequence motifs, and germline gene usage, the majority of cells were located preferentially in clusters 4, 5 and 9 (Figure 4C). Importantly, GP-specific clones from the GFAP:GP and aCD20-GFAP:GP groups were located in clusters 5 and 9, despite the relatively few cells occupying this cluster for the former group. The results of this functional validation strongly suggest that antigen-mediated reactivation of ASCs in the brain are common in both GFAP:GP and aCD20-GFAP:GP groups, but the high number of circulating B cells entering the CNS masked the detection of these cells in the former group. This was in contrast to the antigen-specific cells of STOP:GP and MOG:GP mice, where cells were preferentially located in cluster 4 (Figure 4C, S9C), which corresponds to an ASC-cluster with relatively lower expression of proliferative genes and lower clonal expansion (Figure 2G). Together, this further suggests that B cells in the CNS are capable of expansion in response to reencounters with cognate neo-self antigen.

**Figure 4.**
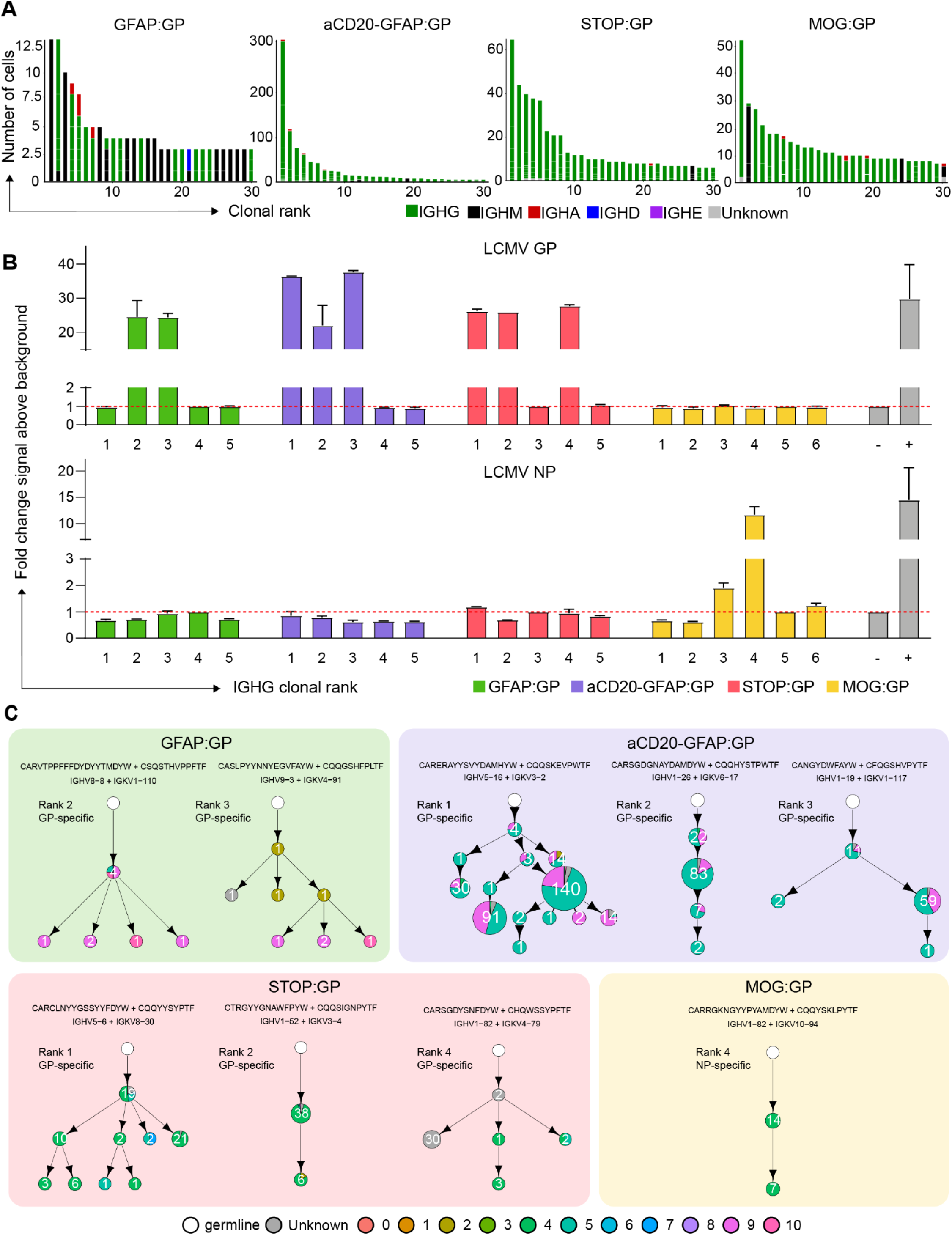
The antigen specificity of clonally expanded and class-switched ASCs of the brain. A. The relationship between the number of cells per clone and the number of clonally related antibody variants within the indicated clone. The thirty most expanded clones were selected per experimental group. Clone was determined by grouping those B cells containing identical CDRH3+CDRL3 amino acid sequences. Variants within each clone are separated by a white line. Bar color refers to the isotype corresponding to the highest fraction of cells within the variant. B. ELISA signal against lymphocytic choriomeningitis virus (LCMV) glycoprotein (GP) and nucleocapsid protein (NP). Clonal rank was determined within each group based on the highest number of cells within each clonotype. C. Mutational networks of those specific clones. Nodes represent unique antibody variants (combined variable heavy [VH] and variable light [VL] chain nucleotide sequence) and edges demonstrate sequences with the smallest separation calculated by edit distance. Nodes are colored by transcriptional clusters. The size and label of the nodes indicate how many cells express each full-length antibody variant. Clone was determined by grouping those B cells containing identical CDRH3+CDRL3 amino acid sequences. Only cells containing exactly one VH and VL were considered. The germline node represents the unmutated reference sequence determined by 10X Genomics cellranger. CDR3 Sequence motifs on top of each network and corresponding V genes. Color corresponds to biophysical properties.

Given that we only detected LCMV NP-specificity amongst the most expanded clones from the MOG:GP mice (Figure 4B), we questioned whether the constitutive expression of GP by oligodendrocytes resulted in a deletion of GP specific B cells and thereby to B cell tolerance in the CNS. In contrast to brain-infiltrating B cells, ELISA on serum from infected MOG:GP mice (50 dpi) demonstrated reactivity to LCMV GP and LCMV NP (Figure 5A), which was in accordance with previous results that MOG:GP mice did not induce peripheral tolerance in secondary lymphoid organs (Page et al. 2021) and suggesting a tolerance specifically within the CNS. Given a recent report of local B cell tolerance to CNS antigens in the meninges (Wang et al. 2021), we profiled deeper into the expanded repertoire to uncover GP-specificity and investigated tolerance in secondary lymphoid organs occurring in our model. We therefore recombinantly produced an additional 10 clones from the expanded repertoire in the CNS of MOG:GP mice, which uncovered additional NP- but not GP-specific B cells (Figure 5B). To provide further support for local CNS tolerance, we calculated the number of GP-specific clones (as determined in the other three experimental groups) present throughout the MOG:GP repertoire. Although minor clonal overlap was detected across all groups for both IgM/IgD and IgG isotypes when disregarding specificity (Figure 5C), we did not detect any CDR3 sequences in the MOG:GP mice that corresponded to previously discovered GP-specific clones (Figure 5D). Taken together, these findings support a model in which a constitutive expression of LCMV GP by glial cells induces localized elimination of self-reactive B cells and thus CNS tolerance.

**Figure 5.**
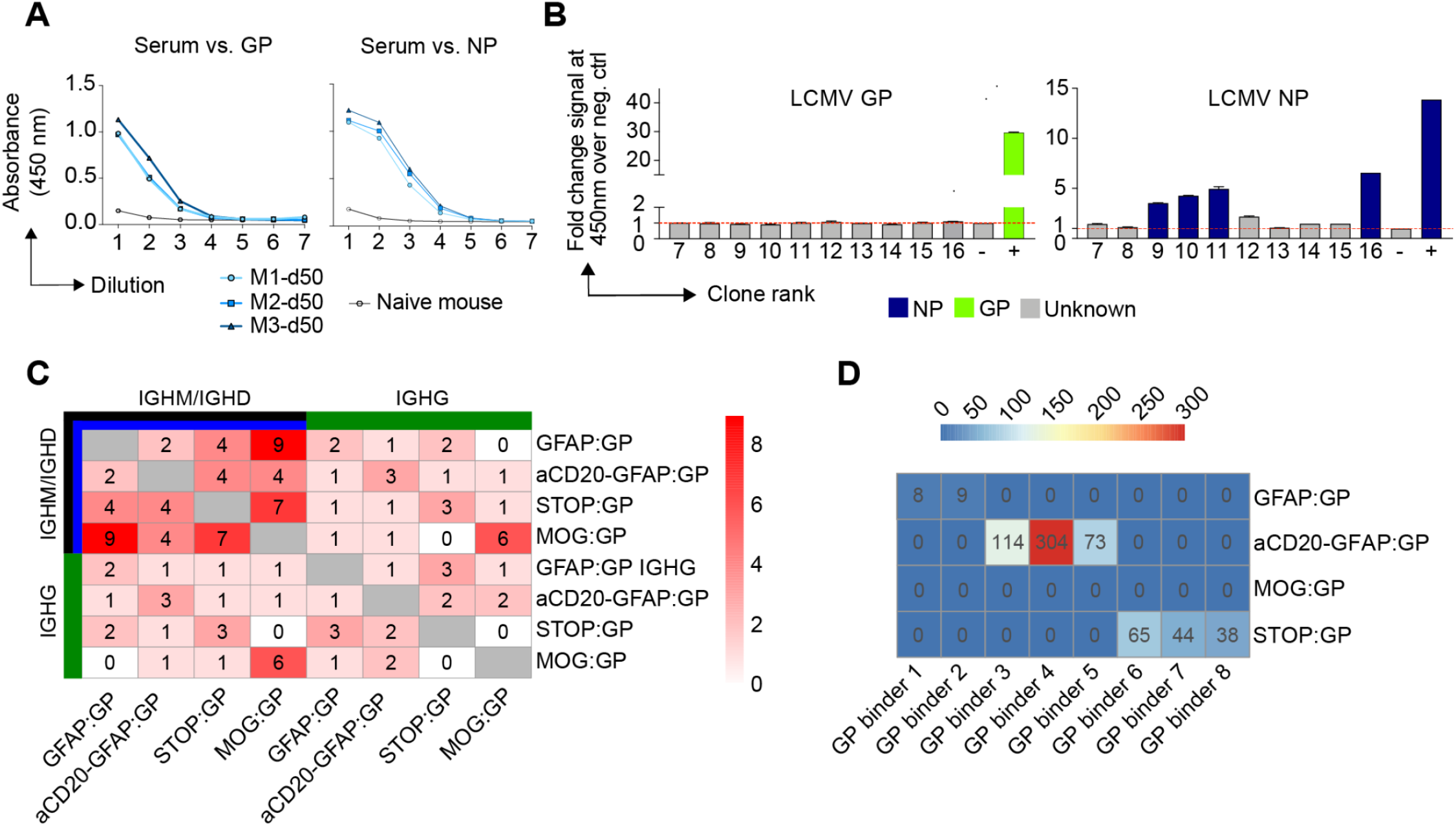
Investigation of CNS-mediated local tolerance. A. ELISA on 5-fold pre-diluted serum (1:100) from uninfected mouse (naive) and infected MOG:GP mice (50 dpi) tested against lymphocytic choriomeningitis virus (LCMV) glycoprotein (GP) and nucleocapsid protein (NP). B. ELISA signal against LCMV GP and LCMV NP for an additional 10 clones from the expanded repertoire of MOG:GP mice (Figure 4A). C. Number of IgM or IgG clones found in more than one repertoire. Clonotyping was performed based on those B cells containing identical HCDR3 + LCDR3 amino acid sequences D. Shared CDR3 amino acid GP-specific sequences across all conditions.

### Proliferating and class-switched ASCs are present across the brain and spinal cord following infection and induction of neo-self antigen

After confirming phenotypic and functional properties of CNS B cells, we next questioned where proliferating and class-switched ASCs were located within the CNS. We detected CD138+ IgG+ cells within the brains of all experimental groups that were not present in naive mice (Figure 6A). CD138+ IgG+ cells co-expressing Ki67 could be detected in various anatomical regions of aCD20-GFAP:GP and GFAP:GP mice such as the meninges, the perivascular space and the parenchyma (Figure 6B). Notably, we detected several instances of CD138+ IgG+ Ki67+ cells located in clusters (Figure 6C), in line with our hypothesis of antigen-mediated proliferation. In contrast, CD138+ IgG+ cells from STOP:GP mice were almost entirely detected in the meninges and rarely in the perivascular space or the parenchyma. Finally, we observed several instances where two CD138+ IgG+ cells appeared to maintain membrane connections (Figure 6C), suggesting that cell division occurs locally within the CNS (Figure 6C). Interestingly, we additionally observed that while the connected cells maintained expression of CD138 and IgG, only one of the two cells had detectable expression of KI67 (Figure 6C), which may indicate that ASCs undergo asymmetric cell division in the CNS. Taken together, our histological data supported our hypothesis that virus-specific B cells can be activated by encountering neo-self antigen in the CNS and adopt proliferative profiles.

**Figure 6.**
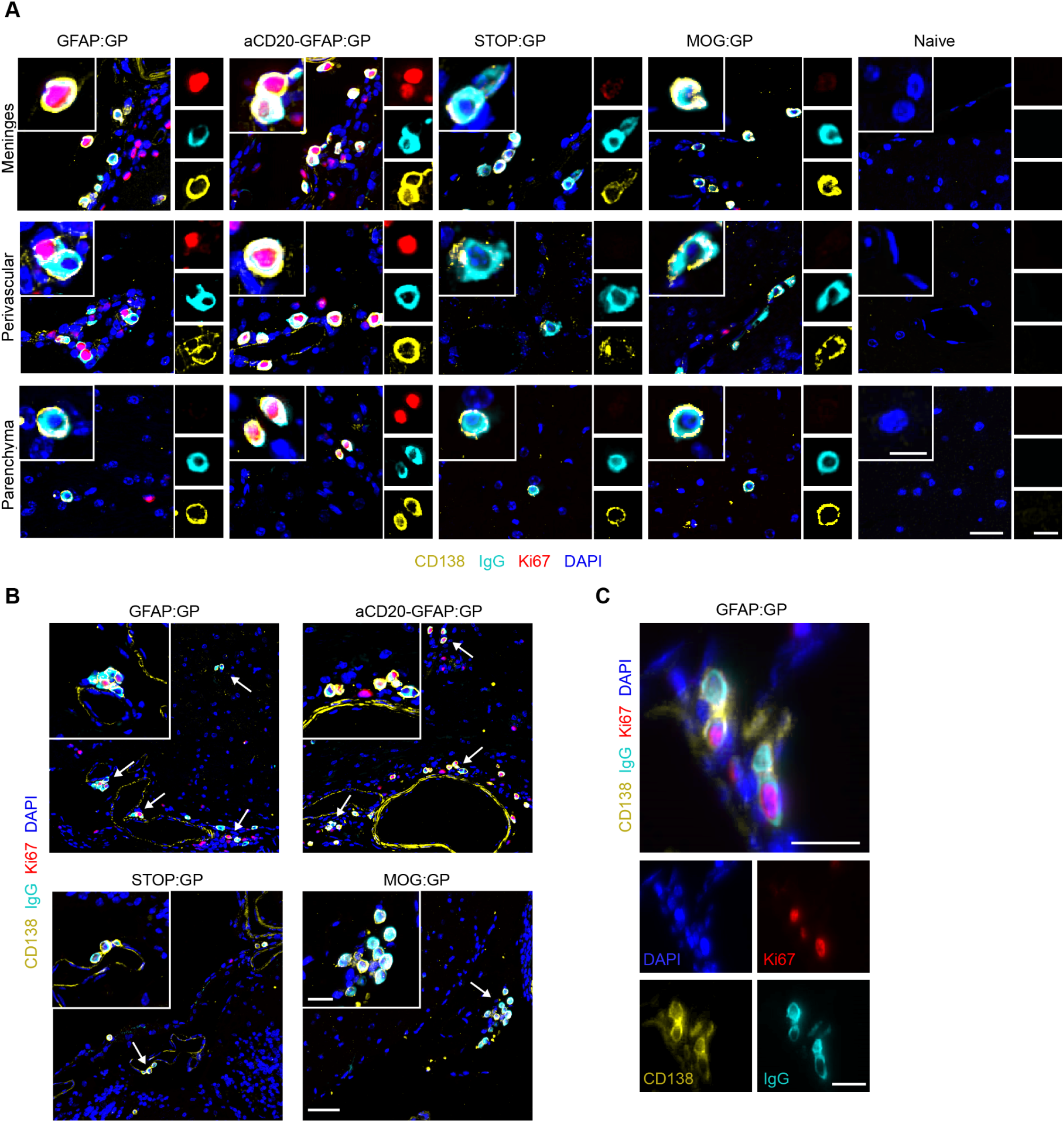
Representative immunostainings for CD138, IgG, Ki67 and DAPI in brain sections of indicated groups. A. CD138+ IgG+ Ki67+/- are mostly located in meninges, perivascular space and parenchyma. Scale: 25um. Inset: 10um. B. CD138+ IgG+ Ki67 cells often localize in clusters (arrows). Scale: 50um. Inset: 20um. C. CD138+ IgG+ Ki67+ cells under cell division. Scale 20um. One representative of at least two independent experiments is shown for A-C.

### Clonally expanded, class-switched, ASCs detected in the cerebrospinal fluid of human neurological diseases

We next questioned whether our experimental findings could be recapitulated in patients affected with inflammatory CNS conditions (Figure 7A). We therefore performed single-cell immune repertoire sequencing of the cerebrospinal fluid from three relapsing multiple sclerosis patients. After filtering out T cells based on expression of CD3e, CD4 and CD8a expression, we performed unsupervised clustering and subsequently visualized the transcriptional landscape. This demonstrated the presence of multiple distinct B cell phenotypes shared across multiple sclerosis (MS) patients (Figure 7B). Consistent with our murine model, a distinct population of cerebrospinal fluid (CSF) B cells could be distinguished on the UMAP by ASC-associated genes, such as *SDC1 (CD138), TNFRSF99 (TACI), SLAMF7*, and *PRDM1* (*BLIMP1*) (Figures 7C, 7D). Of the two major ASC clusters (clusters 0 and 4), the latter also expressed the proliferative marker MKI67 (Figure 7C). We next integrated immune receptor information onto the transcriptional landscape and observed that, in line with our murine model, the ASC clusters coincided with IgG expression (Figure 7E). As we had discovered that clonally expanded, IgG-expressing ASCs were antigen-specific, we next questioned whether comparable levels of clonal expansion could be detected in the CSF of MS patients. Overlaying expansion information for those clones supported by at least two distinct cell barcodes revealed that the vast majority of expanded clones were localized in the IgG-expressing ASC population (Figure 7F). Given the high proportion of expanded clones located across all repertoires, we wondered whether somatic variants could be detected amongst the most expanded clones, similar as previously observed for the expanded IgG-expressing ASCs in our murine models. We therefore inferred phylogenetic networks for the most expanded clones (Figure 7G), which indeed confirmed the presence of clonally-related B cells expressing distinct antibodies within the CSF of all patients (Figure 7H). Taken together, these data suggest that the B-cell transcriptional properties in the CSF of MS patients with relapsing disease share properties similar to those observed in our animal models.

**Figure 7.**
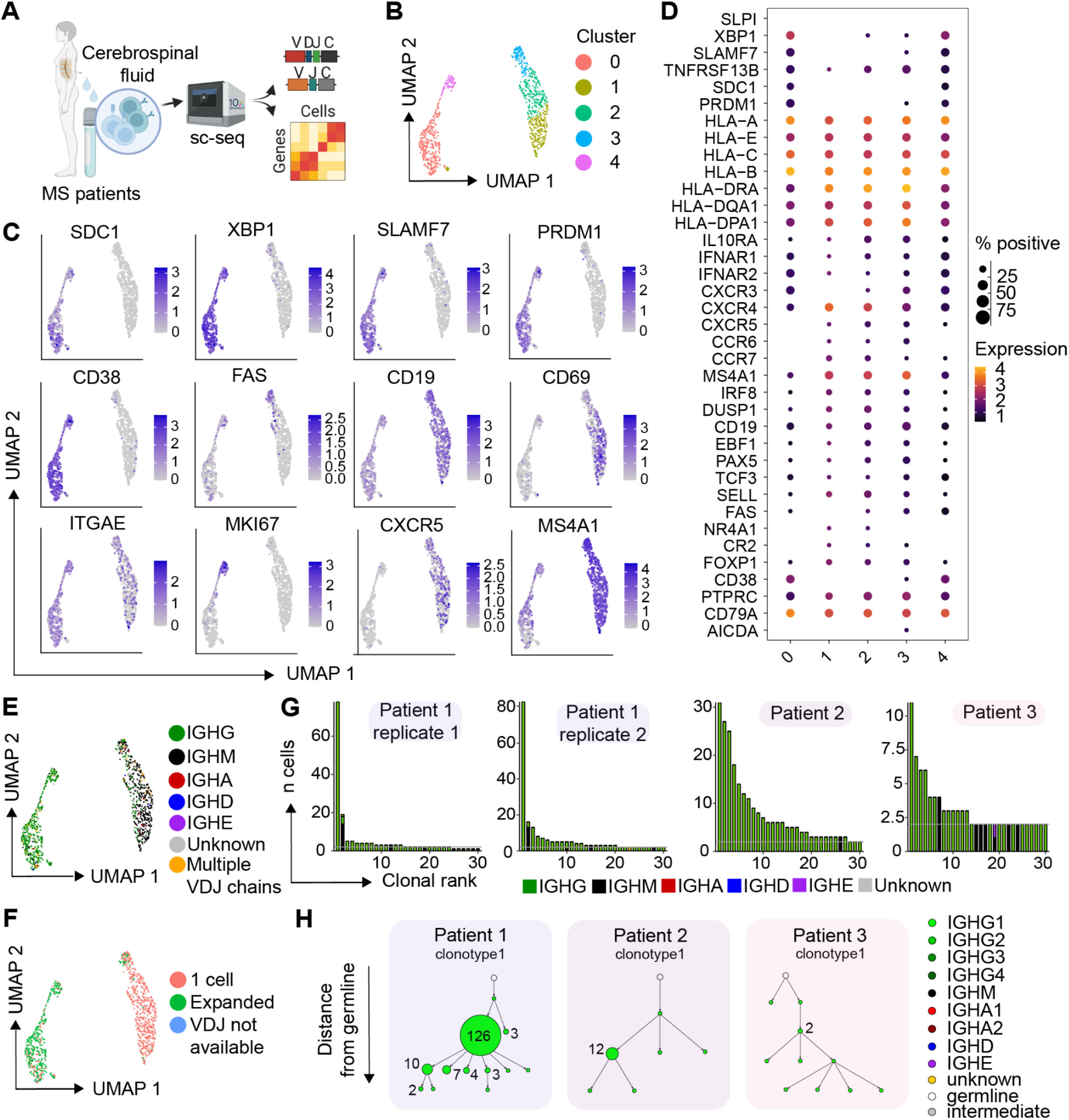
Clonally expanded, class-switched, ASCs detected in the cerebrospinal fluid of multiple sclerosis (MS) patients. A. Schematic overview for the analysis of B cells from MS patients. B. Uniform manifold approximate projection (UMAP) of human MS patients colored by transcriptional cluster membership. C. Gene expression of select genes UMAPs showing gene expression of selected B-cell genes. D. Normalized expression of B-cell markers separated by cluster membership. E. UMAP of human MS patients colored by Isotype. F. UMAP of human MS patients colored by cell expansion. Expanded corresponds to those clones supported by more than one unique cell barcode. G. Clonal frequencies for the 30 most expanded clones per experimental condition. Clones were determined according to 10x Genomics Cell Ranger’s default clonotyping strategy. Clones containing 2 cells are highlighted with a horizontal gray line. Color corresponds to isotype as determined in the VDJ sequencing library. H. Mutational network of the two most expanded clones per patient. Nodes represent unique antibody variants (combined variable heavy chain [VH] + variable light chain [VL] nucleotide sequences) and edges separate those sequences with the smallest edit distance. Node color corresponds to isotype identity. The size and label of the nodes indicate the number of cells expressing a single full-length antibody variant. Clones were determined according to 10x Genomics Cell Ranger’s default clonotyping strategy and only those cells containing exactly one VH and VL chain were included. The germline node represents the unmutated reference sequence determined by 10x Genomics Cell Ranger.

## Discussion

B cells and their corresponding antibodies have been implicated in various chronic autoimmune disorders of the CNS. Here, we leveraged well-defined murine models of autoimmunity and infection to discover phenotypically and functionally diverse populations of B cells in the CNS. We discovered that clonally expanded and IgG-expressing populations of virus-specific ASCs persist in the CNS weeks after viral clearance. Furthermore, we show that these ASCs in the CNS reside in niches, but respond and proliferate following exposure to local autoantigens. Interestingly, this observation of IgG-expressing is in contrast to prior studies that have demonstrated IgA-secreting plasma cells in the meninges during homeostasis (Fitzpatrick et al. 2020). Similar to the experimental models, we observed a population of class-switched, clonally expanded, somatically hypermutated, and proliferating ASCs in the CSF of relapsing MS patients.

Consistent with the recently highlighted link between viral infection and MS (Bjornevik et al. 2022), our data show that this ASC population described here represents expanded cells within the CNS that are potentially virus-specific and directed against antigens expressed in resident cells of the CNS. This is further supported by the observation that an expanded B cell clone in the CSF and blood of a MS patient was recently shown to be cross-reactive against both Epstein-Barr virus (EBV) and the glial cell adhesion molecule GlialCAM (Lanz et al. 2022). It is thus likely that the phenotypic properties of this subset can be used to inform antigen specificity of the clonally expanded cells in human CSF in the context of neurological disorders which would substantiate the link between infection and neurological autoimmune disease. B cells are recruited in an antigen-independent manner under inflammatory conditions and can persist without necessarily exhibiting specificity for an antigen within the CNS, complicating the identification of pathophysiologically relevant B cells in inflammatory CNS conditions (Tesfagiorgis et al. 2017). The approach presented here allows for the screening and identification of disease-relevant B cells and can constitute a new avenue for the development of targeted therapies in the context of CNS disorders. In addition, given that B cells efficiently present antigen and can activate cognate T cells (Chen and Jensen 2008), further profiling the specificity of the antigen presenting B cells discovered here could elucidate potential disease-relevant MHCII epitopes that would be presented to T cells.

Of note this subset of murine ASCs was entirely absent in the CNS of naive mice and in the widely used MOG_35-55_- and rMOG_1-125_-induced EAE mouse models (Yermanos, Neumeier, et al. 2021; Shlesinger et al. 2022). While minor clonal expansion of B cells was present in both naive and EAE conditions, the recovered B cells demonstrated either naive or memory B cell phenotypes and entirely lacked ASC gene signatures, class-switching, contemporary clonal variants, and expression of cell cycle and proliferation genes in contrast to MS patients as found in this study. Thus, our recently developed murine model as harnessed in the current work may not only serve as a valuable tool to profile and explore niches of CNS resident memory T cells, but also B cells with relevance to human neurological disease conditions. The formation and maintenance of resident memory B cells have been recently described in the lungs (Tan et al. 2022; Allie et al. 2019; Barker et al. 2021), with certain populations demonstrating ASC phenotypes upon reactivation (MacLean et al. 2022). Our observations that B cell populations persist in the CNS following aCD20 administration, suggests that at least a fraction of these cells may be tissue resident. This is in line with previous findings showing that B cells can be recruited and persist in an antigen-independent manner in the CNS (Tesfagiorgis et al. 2017; Pollok et al. 2017).

One further observation was the lack of B cells in the CNS reactive against LCMV GP when constitutively expressed as a neo-self antigen in oligodendrocytes. This suggests that the constant presence of neo-self antigen in the CNS results in deletional B cell tolerance preventing residence of autoantigen-specific B cells. The idea that deletional tolerance is occurring locally within the CNS and related niches following the recruitment of antigen-specific B cells is further supported by the fact that serum titers against the neo-self antigen remained unchanged. Such local tolerance mechanisms could potentially be related to hematopoietic niches within the meninges and skull that have been described as immune cell reservoirs and instruct B cells during development (Cugurra et al. 2021; Brioschi et al. 2021). Furthermore, it has been demonstrated that negative selection against CNS antigens occurs during B cell development within the meninges, although this investigation was restricted to MOG-specific B cells (Wang et al. 2021). Mechanistically, CSF-mediated cross-talk between the brain, meninges, and skull bone marrow niches may contribute to the distribution and accessibility of the neo-self antigen locally within the CSF. This would be in line with recent findings showing that intra cisterna magna injections drain to local skull bone marrow niches but not those present in the tibia (Mazzitelli et al. 2022), which would further prevent the induction of peripheral tolerance. To what extent and how such tolerance mechanisms become dysfunctional in human chronic autoimmune diseases will be an important task to resolve in future work.

Overall, our study highlights the clonal dynamics of B cells populating the CNS during infection and autoimmunity in experimental models and human disease conditions. It further emphasizes that B cells can undergo clonal selection and respond to autoantigens expressed in the CNS and it delineates the transcriptional characteristics of ASC populations that can be used to determine the antigen origin of B cell responses taken place within the CNS. Finally, our study serves as a starting point to unravel the poorly understood underpinnings of B cell dysregulation in chronic inflammatory CNS diseases.

---

**Figure S1.**
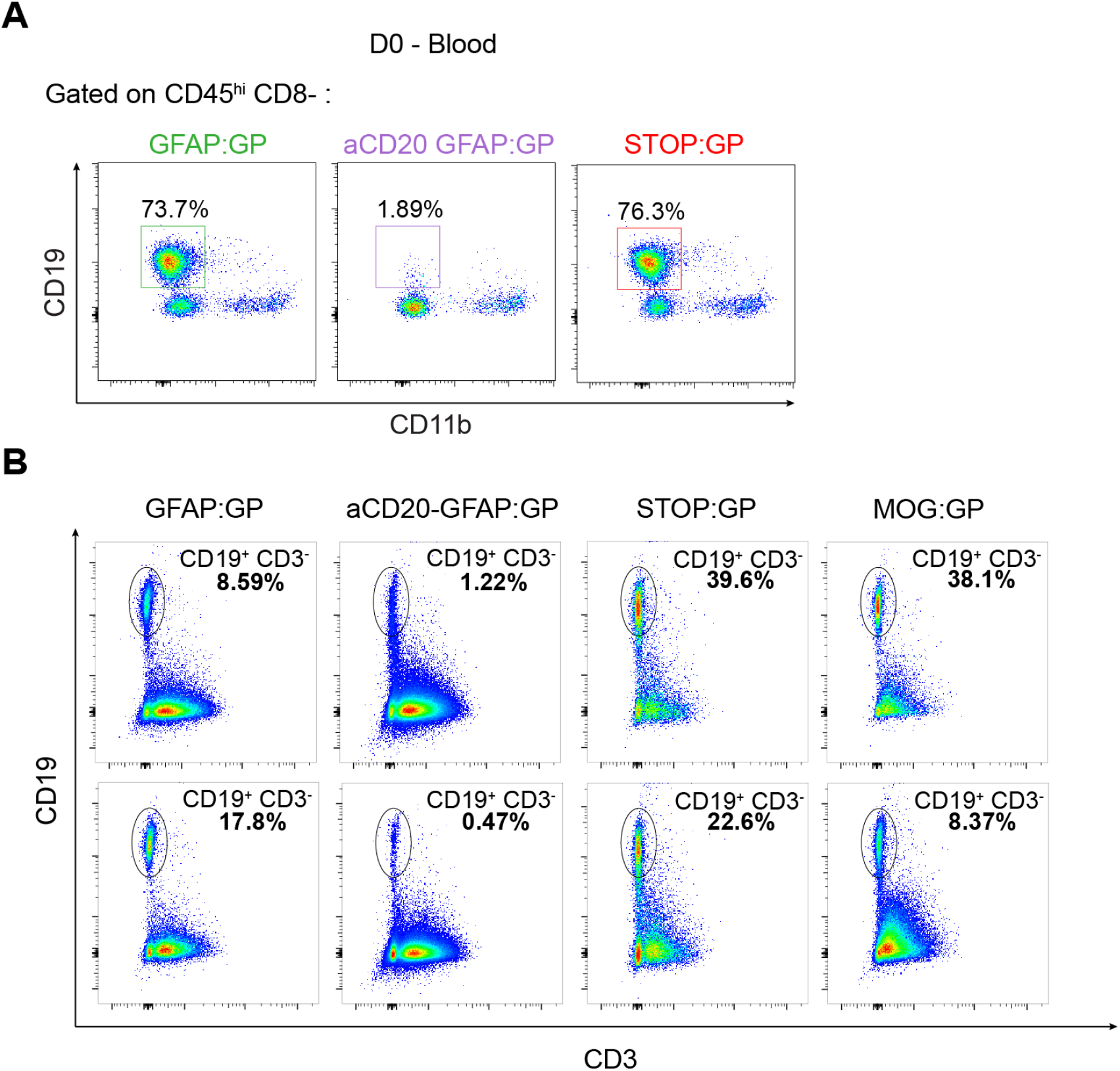
Flow cytometry plots of sorted populations. A. Flow cytometry plots of CD45hi CD8-CD11b CD19+ cells in the blood at day 0 of tamoxifen administration in indicated groups. Numbers represent the percentage of positive cells. B. Flow cytometry dot plot analysis of sorted CD19^+^/CD3^-^ B cell populations per mouse for each experimental group for remaining mice not shown in Figure 1F.

**Figure S2.**
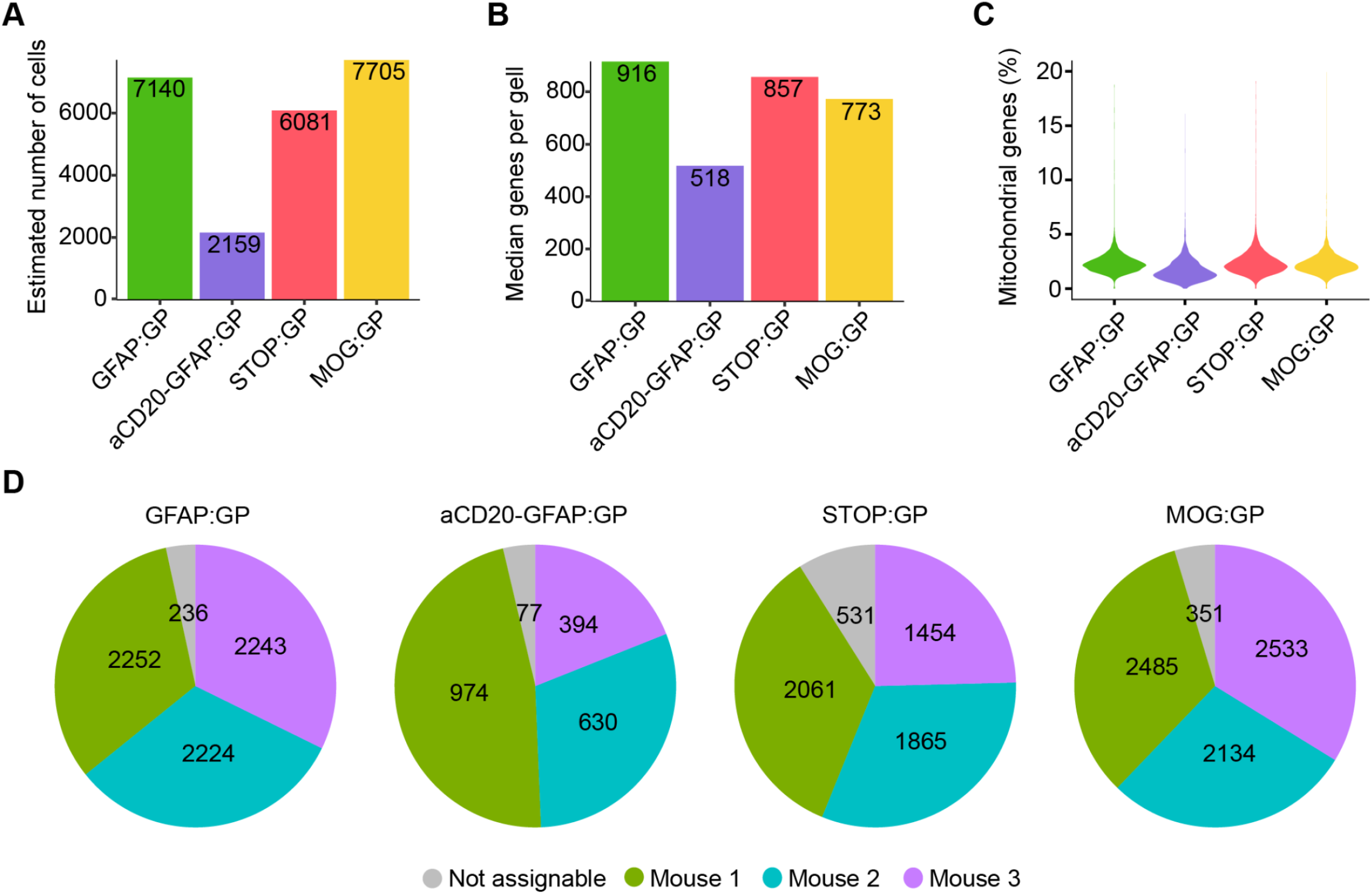
General statistics following single-cell sequencing. A. Number of cell barcodes in gene expression (GEX) sequencing libraries B. Number of unique genes per cell. C. Average number of mitochondrial reads per cell. D. Number of cells per hashing barcodes.

**Figure S3.**
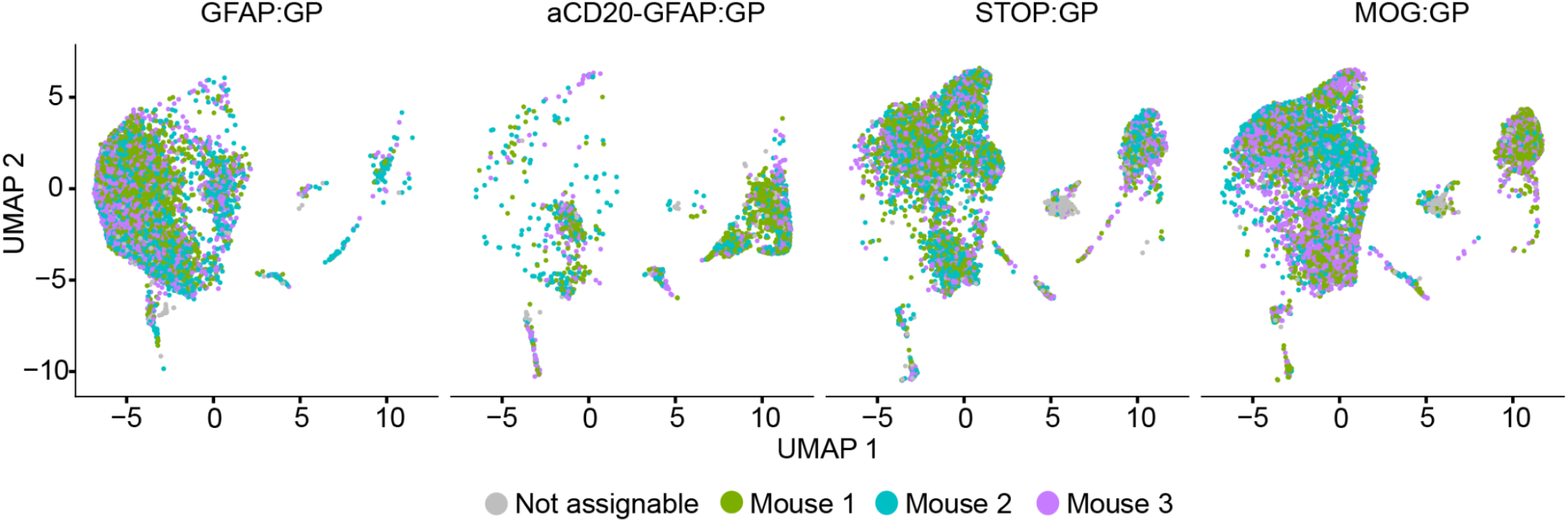
Uniform manifold approximation projection (UMAP) split by sample and colored by mouse-specific DNA barcode demonstrating biological reproducibility.

**Figure S4.**
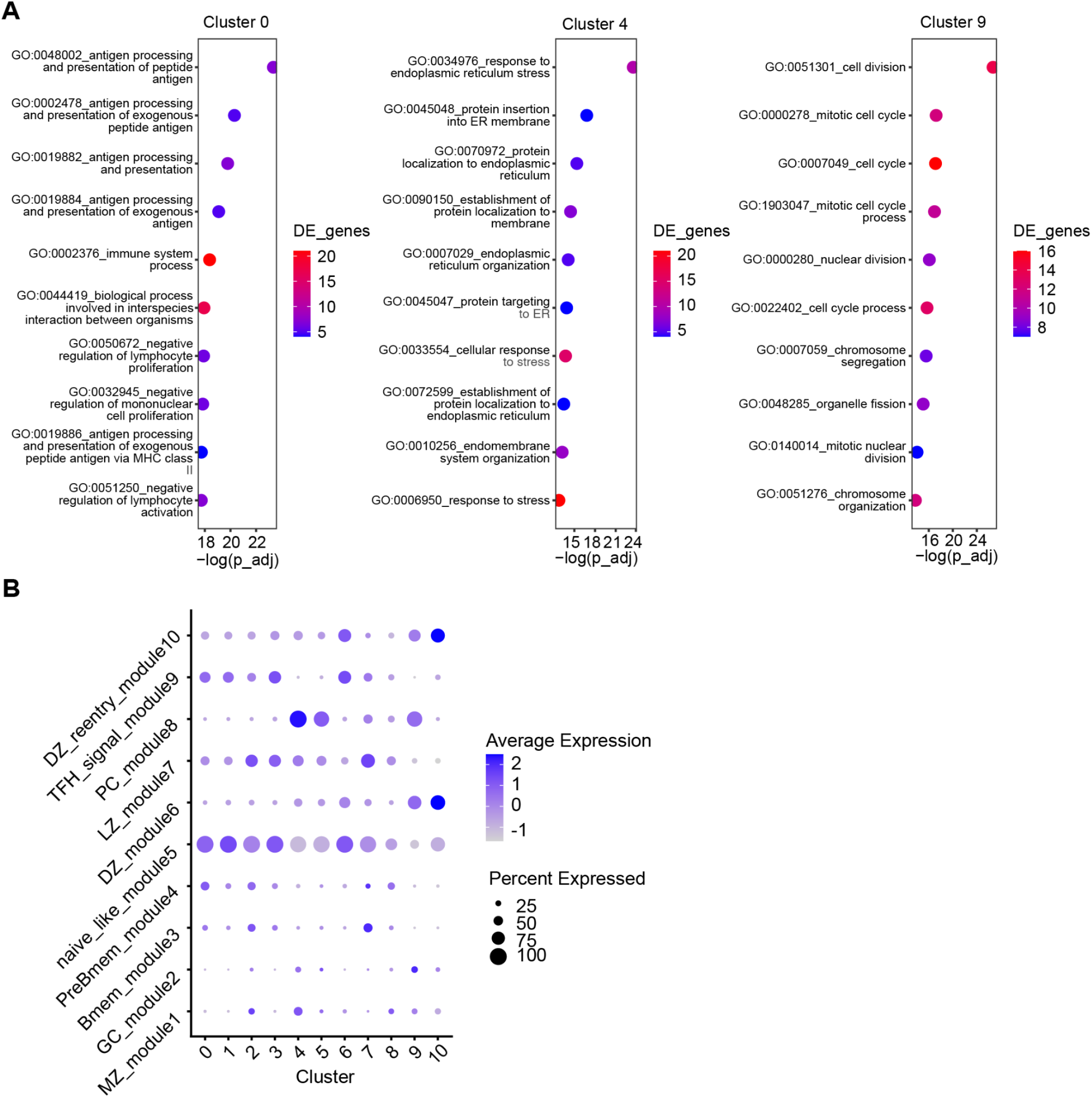
Gene ontology (GO) and gene modules to classify B cell phenotypes. A. GO term enrichment of the 10 most upregulated genes from transcriptional clusters 0, 4 and 9 based on average log fold change. The color of each dot corresponds to the adjusted p-value and the size corresponds to the number of genes. Ratio corresponds to the number of differentially expressed genes relative to the number of total genes corresponding to each GO term. B. Dottile plot showing B cell subset assignment across all clusters based on expression of genes defining B cell phenotypes. The intensity of each dot corresponds to the average expression of all cells within a given cluster and the size corresponds to the percentage of cells with detectable gene expression.

**Figure S5.**
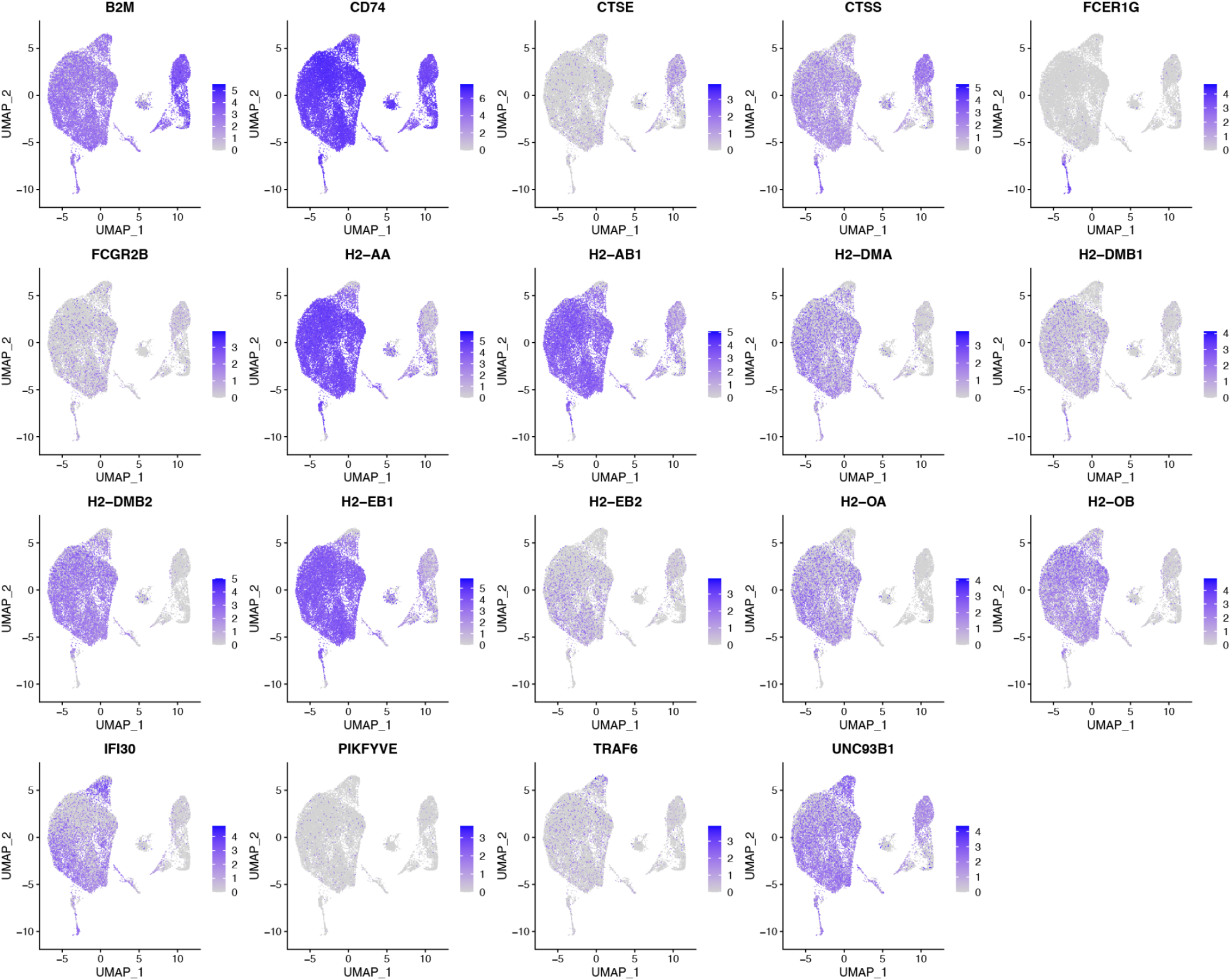
Gene expression of select genes associated with class II antigen presentation.

**Figure S6.**
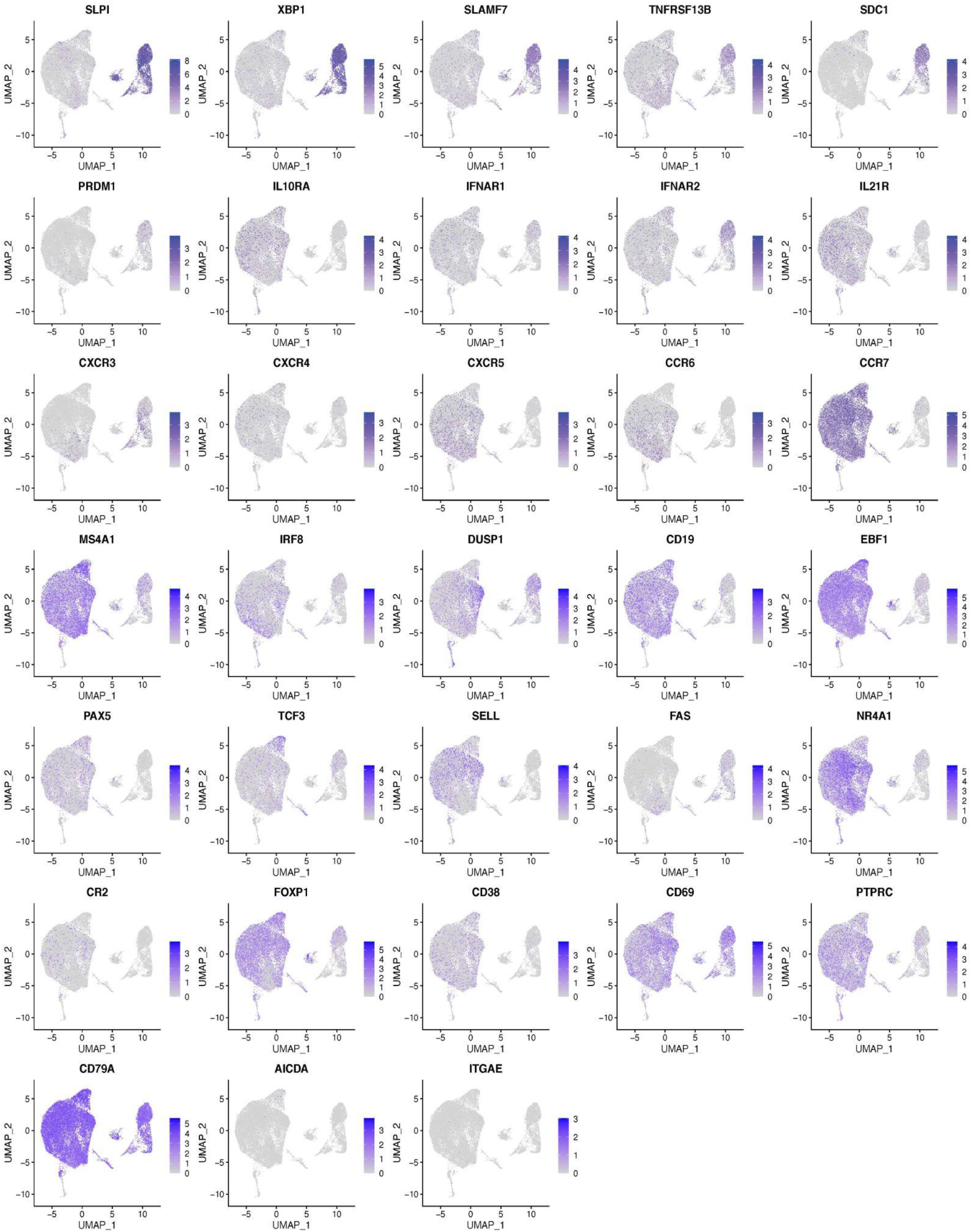
Gene expression of select genes for all cells from all conditions.

**Figure S7.**
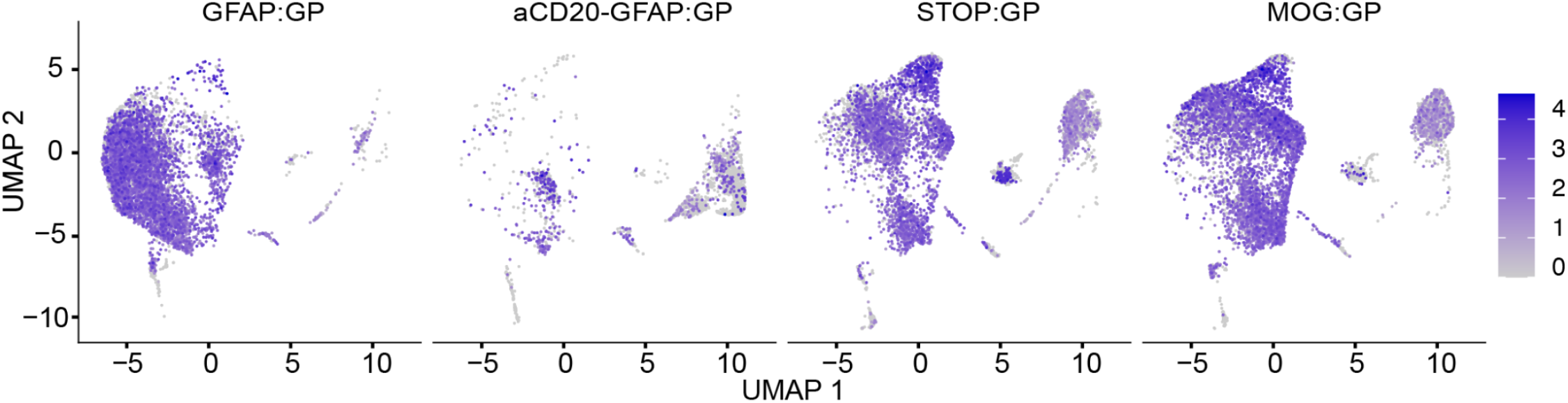
Gene expression of *CD20 (MS4A1*) for all cells from all conditions.

**Figure S8.**
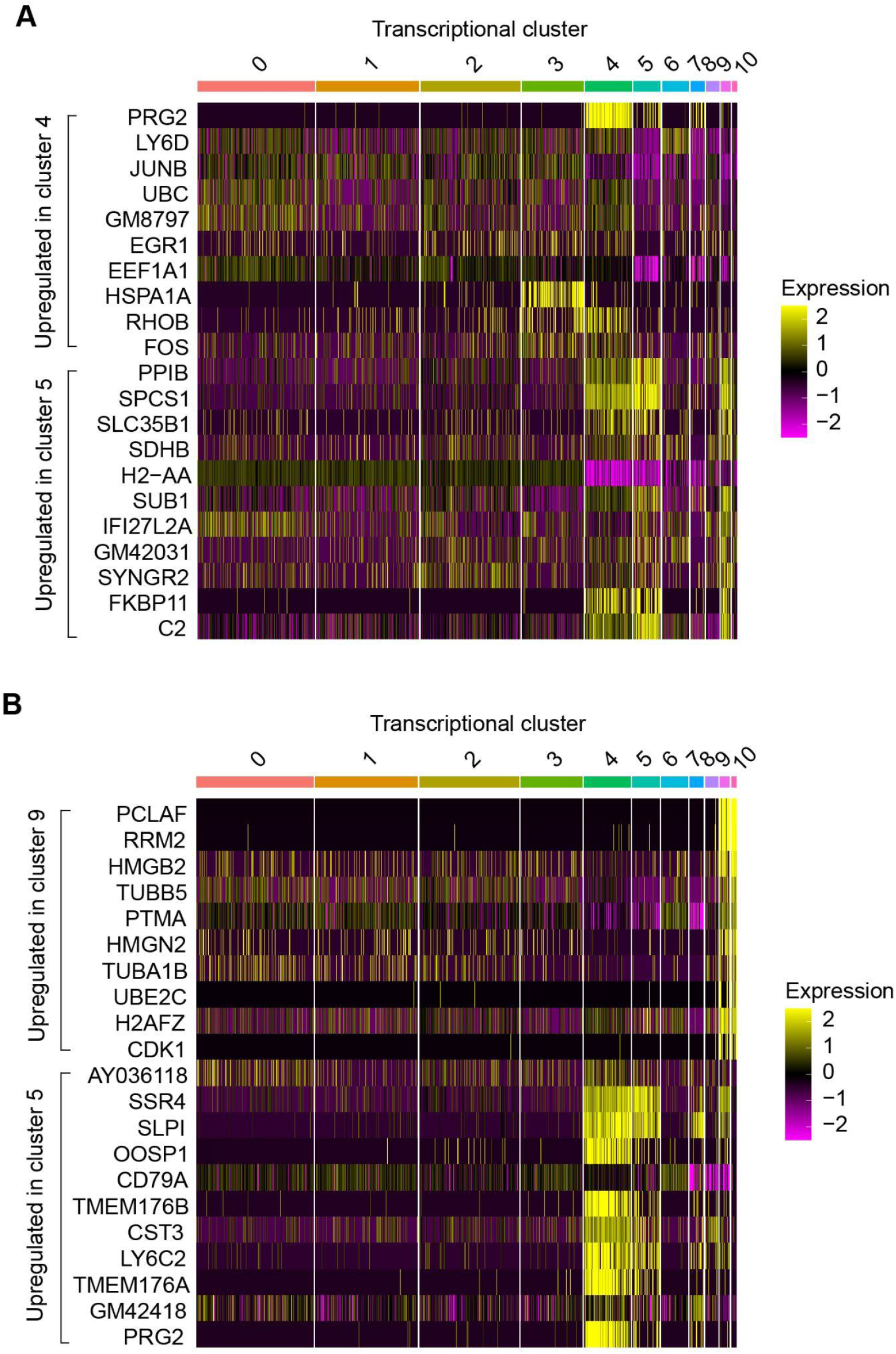
Distinct gene expression signatures of ASCs located in transcriptional clusters 4, 5, and 9. A. Differentially expressed genes between clusters 4 and 5 ranked by average log-fold change. B. Differentially expressed genes between clusters 9 and 5 ranked by average log-fold change.

**Figure S9.**
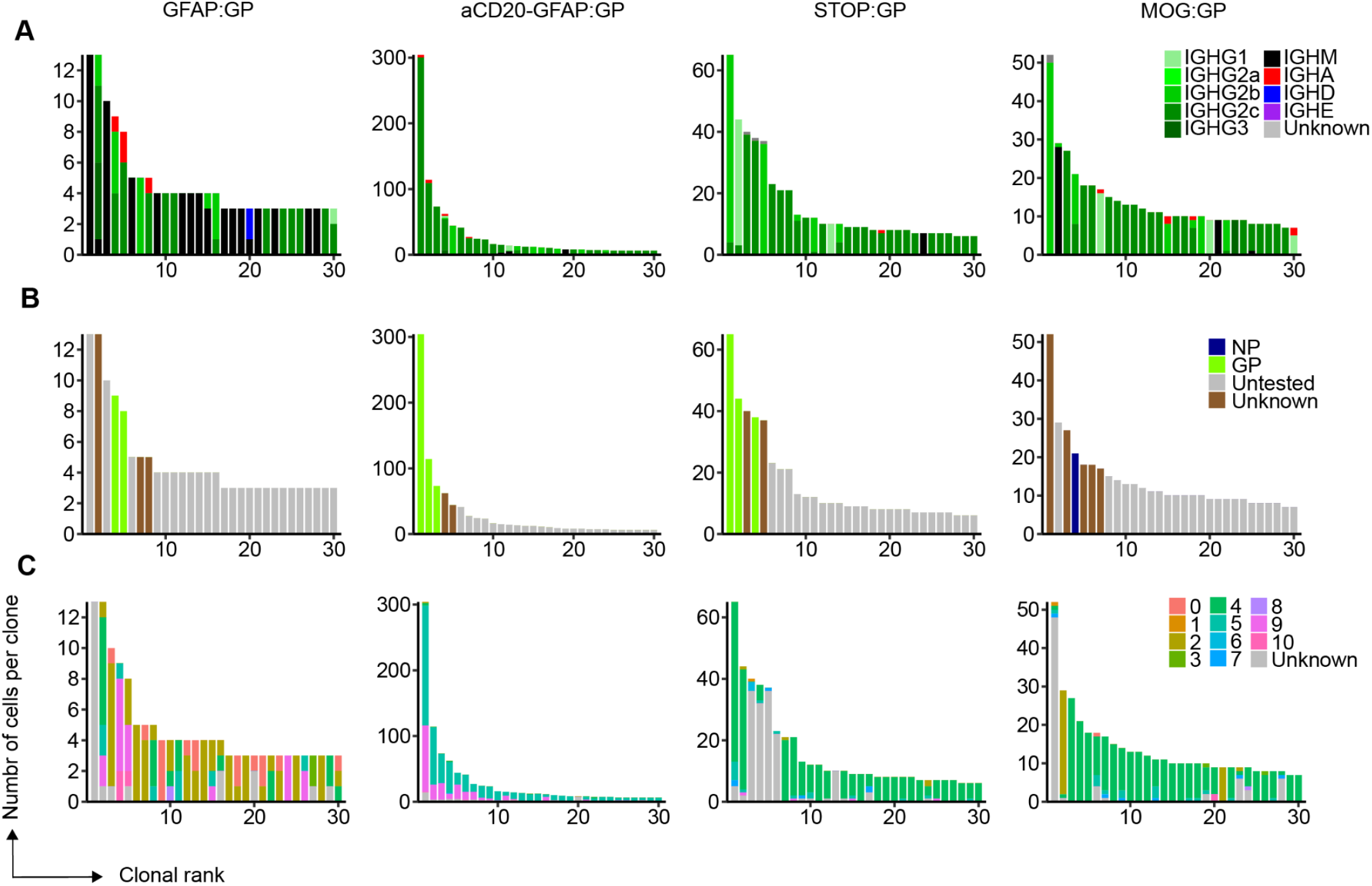
Isotype, specificity, and transcriptional cluster of the 30 most expanded clones per experimental condition. Clonotyping was performed based on those B cells containing identical CDRH3 + CDRL3 amino acid sequences. Cells or clones were colored based on their A. isotype, B. antigen-specificity (one variant tested per clonal family), or C. transcriptional clustering based on gene expression data.

**Figure S10.**
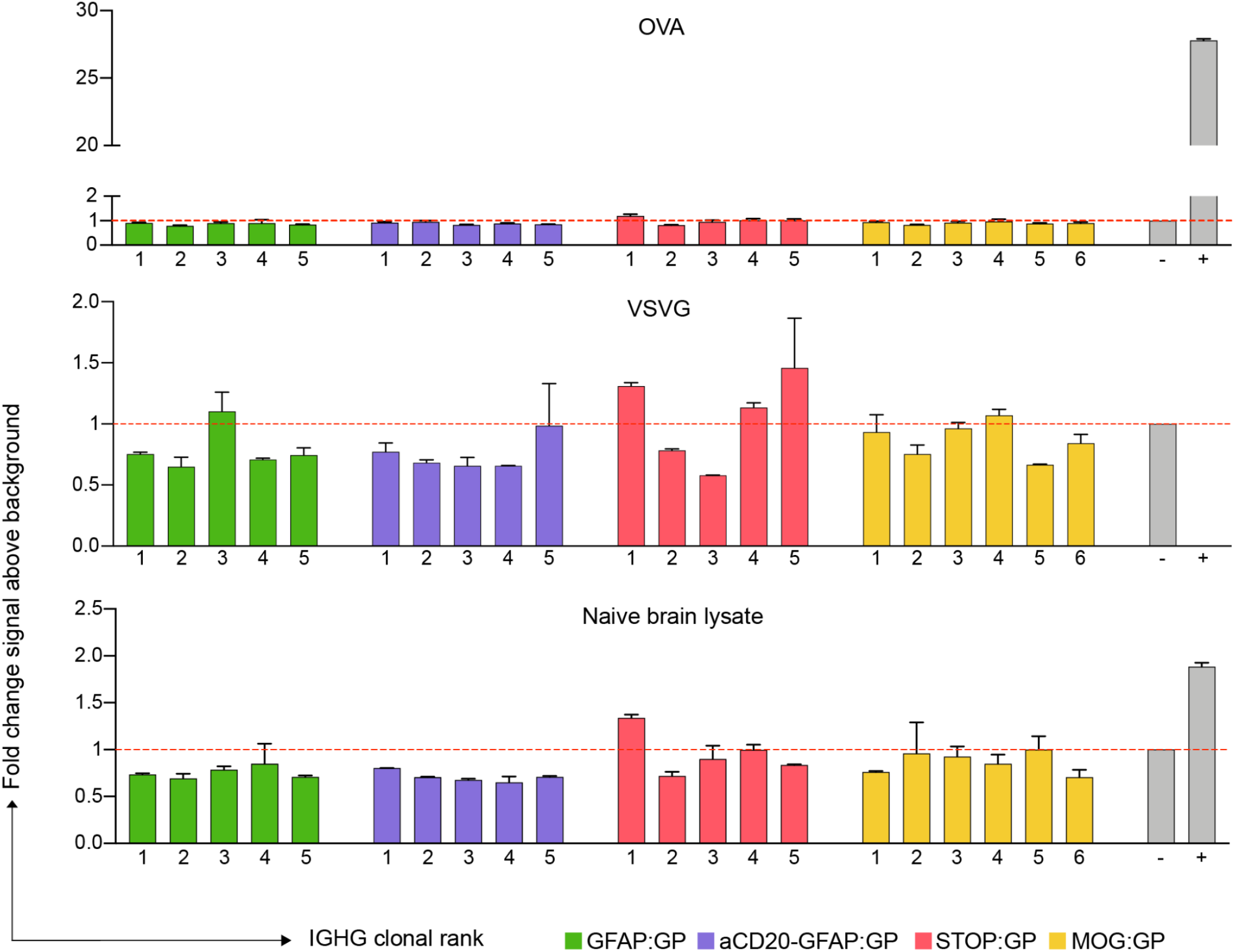
Antigen specificity of clonally expanded and class-switched ASCs. ELISA signal against ovalbumin (OVA), vesicular stomatitis virus glycoprotein (VSVG) and naive brain lysate from naive B6 mouse. Clonal rank was determined within each group based on the highest number of cells within each clonotype.

## Methods

### Mice

The C57Bl/6J GFAP:GP (GFAP-Cre^ERT2 tg/+^;Stop-GP^flox/+^) mouse line was bred by crossing C57Bl/6 GFAP-CreERT2 mice (Hirrlinger et. al 2006) expressing the TAM-inducible Cre-recombinase under the astrocyte-specific GFAP promoter with C57Bl/6J Stop-GP mice (Vincenti et al. 2022). The C57BL/ 6J MOG:GP (MOGi^Cre/+^: Stop-GP^flox/+^) mouse line was generated as described previously (Page et al. 2021). All mice were bred and lodged under specific-pathogen-free P2 conditions in the animal facilities of the University Medical Center of Geneva. Age-matched male and female sex mice between 6 and 12 weeks of age were used for experiments. Animal experiments were authorized by the cantonal veterinary office of Geneva and performed in conformity with institutional guidelines and Swiss federal regulations under license number XGE3920D.

### Virus infections

The recombinant LCMV-GP58 were generated and propagated on baby hamster kidney 21 cells BHK21 (ATCC^®^ CCL-10) (Flatz et al. 2006). Virus stocks were titrated using MC57G cells (ATCC^®^ CRL-2295) (Kallert et al. 2017). In the recombinant LCMV-GP58, the LCMV full-length-GP was replaced by a fusion protein of the LCMV GP signal sequence including the GP_33-41_ epitope and the vesicular stomatitis virus glycoprotein (VSVG). For transient infection of the CNS, mice were intracranially injected with 10^4^ plaque-forming units (PFU) of rLCMV-GP58 diluted in 30 μl of minimum essential medium (MEM, Gibco).

### Tamoxifen (TAM) administration

TAM (Sigma, T5648) was dissolved in a 9:1 mix of sunflower oil and ethanol to a concentration of 10 mg/ml by mixing the solution for 20min at 50 °C. A dose of 100 μl of TAM solution was administered intraperitoneally twice per day for 4 consecutive days.

### Antibody-mediated depletion

Circulating CD20^+^ cells were depleted by intraperitoneal administration of 200 μg of rat-anti-mouse-CD20-depleting antibody (Biolegend, 152104) two days before the start of TAM administration. Depletion of circulating CD20+ B cells was confirmed by flow cytometry.

### Rotarod

The rotarod test was used to monitor the ability of motor coordination and balance of mice. Mice were placed on a rotating rod (Rotarod 7650; Ugo Basile Biological Research) constantly accelerating from 10 to 80 rounds per minute for a maximum of 180 s. Endurance time was monitored, and the two best runs out of three at each time point were averaged for analysis. Animals were habituated and trained to the rotarod daily from day −3 to day 0. Displayed values (% latency to fall) are normalized to the running values of mice at day 0.

### Cell isolation and flow cytometry

For the isolation of cells in the CNS, mice were anesthetized and transcardially perfused with PBS. CNS tissues were cut and digested in DMEM with Collagenase A (1 mg/ml, Roche) and DNase I (0.1 mg/ml, Roche) at 37°C for one hour. Cells were homogenized using a 70-μm cell strainer (BD Biosciences) before leukocytes were separated using 30 and 70% Percoll gradients and subsequently stained for 30 minutes in FACS buffer (PBS/2.5%FCS/10mM EDTA/0.01% NaN3) with the following primary antibodies: CD45-PacificBlue (Biolegend, 103126), CD19-biotin (Biolegend, 115503), CD3E-APC (Biolegend, 100311). Oligonucleotide barcodes conjugated to phycoerythrin (PE) and streptavidin (TotalSeq™-C0951 PE Streptavidin, TotalSeq™-C0952 PE Streptavidin and TotalSeq™-C0953) were added to biotinylated anti-CD19 antibodies to maintain sample identity following pooling of B cells for each experimental condition. CD19+ B cells were sorted individually for each mouse using a BD LSRFortessa (BD Biosciences) and BD FACSDiva (BD Biosciences, v8.0.2) using appropriate filter sets and compensation controls. B cells arising from the same experimental condition were subsequently pooled and used as input to single-cell immune repertoire sequencing. Data was analyzed using FlowJo software (v10).

### Single-cell immune repertoire sequencing of murine B cells

For single-cell sequencing, brain B cells (CD19+ CD3-CD45.1+) were FACS-sorted from STOP:GP, aCD8-GFAP:GP and GFAP:GP mice 7 days after start of TAM, and from MOG:GP mice 47 days after i.c infection. Combined single-cell transcriptome and immune repertoire sequencing was performed using the 10x Genomics Chromium Single Cell V(D)J Reagents Kit (CG000166 Rev A) according to established methods (Neumeier et al. 2021). Single cells together with gel beads (10x Genomics, 1000006) were encapsulated in gel emulsion microdroplets using 4 lanes of one Chromium Single Cell A Chip (10x Genomics, 1000009) with a target loading of 13,000 cells per reaction. Subsequent cDNA amplification was performed using 14 cycles. The samples were then split for separate GEX, VDJ and feature barcode library preparation (10x Genomics, 1000080). GEX and feature barcode libraries were amplified using the Chromium Single Cell 5’ Library Kit (10x Genomics, 1000006) and BCR libraries were amplified using the Chromium Single Cell V(D)J Enrichment Kit, Mouse B Cell (10x Genomics, 1000072). The final libraries were pooled and sequenced using the Illumina NovaSeq S1 platform at a concentration of 1.8 pM with 5% PhiX.

### Single-cell immune repertoire sequencing of B cells from CSF of MS patients

For single-cell sequencing of B cells in the CSF of MS patients, fresh CSF samples were processed within one hour after collection. CSF samples (3-6 ml) were centrifuged at 300 g for 10 minutes. The pellet was then transferred to a 2-ml tube and further processed using the 10x Chromium Next GEM Single Cell VDJ v1.1 Reagents Kit (CG000208 Rev B) as previously described (https://doi.org/10.1038/s41590-021-00948-8). GEX and BCR libraries were sequenced on an Illumina NovaSeq6000 S4 using read lengths of 150 bp read 1, 8 bp i7 index, 150 bp read 2. The study was approved by the local ethics committee of the Ludwig-Maximilian University Munich (project no. 163-16 and 18-419), and written informed consent was granted by all participants included in the study.

### Immune repertoire analysis

Raw paired-end sequencing files of the GEX, V(D)J, and feature barcode libraries were aligned to the murine reference genome (mm10), V(D)J germlines (GRCm38) and reference feature barcode sequences using 10x Genomics Cell Ranger (v6.0.0). Accordingly, GEX and V(D)J libraries from human samples were aligned to the human reference genome and V(D)J germlines (GRCh38) Specifically, the count argument of Cell Ranger was used to align GEX and feature barcode libraries separately while the V(D)J was aligned using the vdj argument. The filtered feature matrix GEX and V(D)J files as well as unfiltered feature barcode files were supplied into the VDJ_GEX_matrix function of the R package Platypus (v3.1) (Yermanos et al. 2021), which relies on the R package Seurat (v4.0.3) (Satija et al. 2015) for gene expression analysis. For human samples, cells were filtered based on CD3E, CD4 and CD8A expression to exclude T cells from the analysis. Annotations from GEX were transferred to VDJ and vice versa by matching cellular 10x barcodes. Cells containing more than 5% mitochondrial genes were removed from transcriptome analysis. Gene expression was log-normalized with a scaling factor of 10,000 and the mean expression and variance were additionally scaled to 0 and 1, respectively. 2,000 variable features were supplied as input to principal component analysis (PCA) using the “vst” selection method. The first ten dimensions were used to assign the cells to transcriptional clusters with the Seurat functions FindNeighbors and FindClusters (Satija et al., 2015) at a cluster resolution of 0.5 using a graph-based clustering approach incorporating Louvain modularity optimization and hierarchical clustering. UMAP was calculated using the first ten dimensions. Gene expression feature plots and violin plots were created by supplying selected genes to the FeaturePlot and VlnPlot functions of Seurat. The GEX_cluster_genes function from Platypus, which relies on the FindMarkers function from Seurat, was used to calculate differentially expressed genes across clusters and conditions with logfc.threshold set to 0 using the Wilcoxon Rank Sum. Mitochondrial and ribosomal genes were removed when visualizing differentially expressed genes (DEG) using the GEX_volcano and GEX_cluster_genes_heatmap functions from Platypus or supplying the top DE genes as input to GEX_volcano and GEX_gsea functions from Platypus to perform gene ontology and gene set enrichment analyses, respectively. The C7 immunological signatures gene set from the Molecular Signatures Database (MSigDB) was used as an input for the GEX_gsea function, which relies on the R package fgsea (v1.16.0) (Sergushichev 2016). The GEX_GOterm function of Platypus is based on the R package edgeR (v3.14) (Robinson, McCarthy, and Smyth 2010). Clonotyping was performed based on those B cells containing identical CDRH3 + CDRL3 amino acid sequences using the VDJ_clonotype function of the R package Platypus. Only clones containing exactly one VH and one VL sequence were included in the BCR analysis. Heatmpas visualizing the number of public clones and the VH and VL gene usage were created using the R package pheatmap (v1.0.12). For antibodies that were selected for expression in PnP-mRuby hybridoma cells, the full-length VH and VL sequences including framework region 1 to framework region 4 were annotated using MiXCR (v3.0.1) and exported by the VDJRegion gene feature.

### Histology and image analysis

Mice were perfused with PFA, CNS tissue were collected, PFA fixed overnight, paraffin embedded and cut at 2um. For immunofluorescence, endogenous mouse IgG were visualized by incubating tissue sections 1 hour with an AlexaFluor 647-conjugated anti-mouse IgG (JacksonImmunoresearch, 715-605-151). Sections were washed and incubated with Dako REAL peroxidase blocking solution (Dako, K0672) to inactivate endogenous peroxidases and unspecific bindings were blocked (PBS/10% FCS).Tissue sections were incubated with rabbit anti-Ki67 antibody (Abcam, ab66155) overnight at 4°C in Dako REAL antibody diluent (Dako, S2022). To visualized the specific signal, anti-rabbit HRP (Dako, K4003) with amplification (Opal 570, Akoya, FP1488001KT) was used as a secondary system. After wash, sections were incubated with Fab fragment-goat-anti-mouse IgG (JacksonImmunoResearch, 115-007-003) and goat serum to avoid unspecific bindings. Rat anti-CD138 antibody (rat anti-CD138, clone 281-2, Biolegend, 142502) was applied to each section and specific bounds were visualized using a Alexa Fluor488-conjugated anti-rat antibody (JacksonImmunoResearch, 712-545-153). Nuclei were visualized using DAPI (Invitrogen, D1306). For immunohistochemistry, sections were incubated with Dako REAL peroxidase blocking solution (Dako, K0672) to inactivate endogenous peroxidase and Fab fragment-goat-anti-mouse IgG (JacksonImmunoResearch, 115-007-003) to avoid nonspecific bindings. Slides were incubated with a rat anti-B220 antibody (RA3-6B2, eBioscience, 14-0452-85). Bound primary antibodies were visualized with a goat anti-rat HRP (Vector Laboratories, MP-7444-15) and bound secondary antibodies with 3,3’-diaminobenzidine as chromogen (Dako, K3468) and counterstained with Hemalum (Merck, 1.09249.0500) for brightfield microscopy. Coverslips were mounted in Fluoromount aqueous mounting medium (Sigma-Aldrich, F4680) for image acquisition. Immunostained sections were scanned using Pannoramic Digital Slide Scanner 250 FLASHII (3DHISTECH) in 200× magnification. All quantifications were performed manually using Pannoramic Viewer software (3DHISTECH). The examiner was blinded to the experimental group. For representative images, white balance was adjusted, and contrast was enhanced using the tools “levels,” “curves,” “brightness,” and “contrast” in Photoshop CS6 (Adobe). All modifications were acquired uniformly on the entire image.

### Antibody expression and validation

Selected antibodies were produced in PnP-mRuby hybridoma cells as previously described (Parola et al., 2019) and validated using normalized supernatant ELISAs against purified LCMV clone 13 NP, LCMV clone 13 GP, DNP-OVA, insulin (Sigma, I5500), mouse genomic DNA (Sigma, 692339), MOG (Sigma, 692339) as previously described (Neumeier et al. 2021). An anti-mouse IgG-HRP (Sigma, A2554) was employed at 1:1500 and used for detection. Purified antigens were tested at 5 μg/ml while self-made extracts were tested at 100 μg/ml coating concentration. Infected MC57G cells were centrifuged for 10 min at 1600 rpm at 4°C and the pellet was resuspended in 5 ml of PBS. Cells were lysed using a syringe (28G needle), homogenized several times and additionally sonicated (3 times 20 s at 40 MHz) on ice. Anti-mouse beta actin IgG (Sigma, A2228), anti-OVA IgG (in house), anti-NP-IgG PANK1, anti-GP-IgG Wen3.1, anti-insulin IgG E11D7 (Sigma, 05-1066) and anti-dsDNA IgG AE-2 (Sigma, MAB1293) were used as negative and positive controls for respective experiments and employed at 4μg/ml.

### Serum ELISA

ELISA 96 well plates (Corning Incorporated) were coated with purified LCMV NP or LCMV GP at 3 mg/ml, blocked with PBS supplemented with 2% (w/v) milk (AppliChem, A0830) and incubated with 5-fold serial dilutions of 1:100 pre-diluted serum (naive serum served as control). Igk binding was detected using anti-mouse kappa light chain-HRP (Abcam, ab99617) secondary antibody. Binding was quantified using the 1-Step Ultra TMB-ELISA substrate solution (Thermo, 34028) and 1M H2SO4 for reaction termination. Absorbance at 450 nm was recorded on an Infinite 200 PRO (Tecan). All commercial antibodies were used according to manufacturer’s recommendations.

### Data visualization

The graphical abstract and experimental setup were created using Biorender. The R packages ggplot (v3.3.3), ggrepel (v0.9.1), VennDiagram (v1.1.0), Seurat (v1.1.1) and gridExtra (v2.3) were used for data visualization. Donut plots and bar graphs were created using GraphPad Prism^®^ Software version 9. All figures were assembled using Adobe Illustrator for Mac (v26.0.1).

## Data availability

The original contributions presented in this study will be made publicly available on the European Bioinformatics following peer review.

## Acknowledgements

We acknowledge and thank Dr. Christian Beisel, Elodie Burcklen, and Ina Nissen at the ETH Zurich D-BSSE Genomics Facility Basel for excellent support and assistance. We also thank Gregory Schneiter for excellent experimental support.

## Funding

This work was supported by the European Research Council Starting Grant 679403 (to STR), ETH Zurich Research Grants (to STR and AO), an ETH Seed Grant (AY) and support from the “la Caixa” Foundation (ID 100010434, fellowship code: LCF/BQ/EU20/11810041) to MMC. DM is supported by the Swiss National Science Foundation (310030B_201271 & 310030_185321) and the ERC (865026). This study was further supported by the German Research Foundation: DO 420/7-1 (KD and EB), EXC 2145 (SyNergy) – ID 390857198 (LAG, KD and EB), and by the DFG Research Infrastructure NGS_CC (project 407495230) as part of the Next Generation Sequencing Competence Network (project 423957469). NGS was carried out at the Competence Centre for Genomic Analysis (Kiel).

## Competing Interests

There are no competing interests.

## Notes

### Competing Interest Statement

The authors have declared no competing interest.

## References

Agrafiotis, Andreas, Daniel Neumeier, Kai-Lin Hong, Tasnia Chowdhury, Roy Ehling, Raphael Kuhn, Ioana Sandu, et al. 2021. “B Cell Clonal Expansion Is Correlated with Antigen-Specificity in Young but Not Old Mice.” bioRxiv. https://doi.org/10.1101/2021.11.09.467876.

Agrawal, Smriti, Per Anderson, Madeleine Durbeej, Nico van Rooijen, Fredrik Ivars, Ghislain Opdenakker, and Lydia M. Sorokin. 2006. “Dystroglycan Is Selectively Cleaved at the Parenchymal Basement Membrane at Sites of Leukocyte Extravasation in Experimental Autoimmune Encephalomyelitis.” The Journal of Experimental Medicine 203 (4): 1007–19.

Allie, S. Rameeza, John E. Bradley, Uma Mudunuru, Michael D. Schultz, Beth A. Graf, Frances E. Lund, and Troy D. Randall. 2019. “The Establishment of Resident Memory B Cells in the Lung Requires Local Antigen Encounter.” Nature Immunology 20 (1): 97–108.

Allie, S. Rameeza, and Troy D. Randall. 2020. “Resident Memory B Cells.” Viral Immunology 33 (4): 282–93.

Barker, Kimberly A., Neelou S. Etesami, Anukul T. Shenoy, Emad I. Arafa, Carolina Lyon de Ana, Nicole Ms Smith, Ian Mc Martin, et al. 2021. “Lung-Resident Memory B Cells Protect against Bacterial Pneumonia.” The Journal of Clinical Investigation 131 (11). https://doi.org/10.1172/JCI141810.

Barr, Tom A., Ping Shen, Sheila Brown, Vicky Lampropoulou, Toralf Roch, Sarah Lawrie, Boli Fan, et al. 2012. “B Cell Depletion Therapy Ameliorates Autoimmune Disease through Ablation of IL-6-Producing B Cells.” The Journal of Experimental Medicine 209 (5): 1001–10.

Bjornevik, Kjetil, Marianna Cortese, Brian C. Healy, Jens Kuhle, Michael J. Mina, Yumei Leng, Stephen J. Elledge, et al. 2022. “Longitudinal Analysis Reveals High Prevalence of Epstein-Barr Virus Associated with Multiple Sclerosis.” Science 375 (6578): 296–301.

Boronat, Anna, Jeffrey M. Gelfand, Nuria Gresa-Arribas, Hyo-Young Jeong, Michael Walsh, Kirk Roberts, Eugenia Martinez-Hernandez, et al. 2013. “Encephalitis and Antibodies to Dipeptidyl-Peptidase-like Protein-6, a Subunit of Kv4.2 Potassium Channels.” Annals of Neurology 73 (1): 120–28.

Brilot, Fabienne, Russell C. Dale, Rebecca C. Selter, Verena Grummel, Sudhakar Reddy Kalluri, Muhammad Aslam, Verena Busch, Dun Zhou, Sabine Cepok, and Bernhard Hemmer. 2009. “Antibodies to Native Myelin Oligodendrocyte Glycoprotein in Children with Inflammatory Demyelinating Central Nervous System Disease.” Annals of Neurology 66 (6): 833–42.

Brioschi, Simone, Wei-Le Wang, Vincent Peng, Meng Wang, Irina Shchukina, Zev J. Greenberg, Jennifer K. Bando, et al. 2021. “Heterogeneity of Meningeal B Cells Reveals a Lymphopoietic Niche at the CNS Borders.” Science 373 (6553). https://doi.org/10.1126/science.abf9277.

Carbone, Francis R. 2015. “Tissue-Resident Memory T Cells and Fixed Immune Surveillance in Nonlymphoid Organs.” Journal of Immunology 195 (1): 17–22.

Chakravarthi, B. V. S. K., M. T. Goswami, S. S. Pathi, A. D. Robinson, M. Cieślik, D. S. Chandrashekar, S. Agarwal, et al. 2016. “MicroRNA-101 Regulated Transcriptional Modulator SUB1 Plays a Role in Prostate Cancer.” Oncogene 35 (49): 6330–40.

Chen, Xinjian, and Peter E. Jensen. 2008. “The Role of B Lymphocytes as Antigen-Presenting Cells.” Archivum Immunologiae et Therapiae Experimentalis 56 (2): 77–83.

Chitnis, Tanuja, and Howard L. Weiner. 2017. “CNS Inflammation and Neurodegeneration.” The Journal of Clinical Investigation 127 (10): 3577–87.

Clark, Rachael A. 2015. “Resident Memory T Cells in Human Health and Disease.” Science Translational Medicine 7 (269): 269rv1.

Csepregi, Lucia, Roy A. Ehling, Bastian Wagner, and Sai T. Reddy. 2020. “Immune Literacy: Reading, Writing, and Editing Adaptive Immunity.” iScience 23 (9): 101519.

Cugurra, Andrea, Tornike Mamuladze, Justin Rustenhoven, Taitea Dykstra, Giorgi Beroshvili, Zev J. Greenberg, Wendy Baker, et al. 2021. “Skull and Vertebral Bone Marrow Are Myeloid Cell Reservoirs for the Meninges and CNS Parenchyma.” Science 373 (6553). https://doi.org/10.1126/science.abf7844.

Dalakas, Marinos C., Harry Alexopoulos, and Peter J. Spaeth. 2020. “Complement in Neurological Disorders and Emerging Complement-Targeted Therapeutics.” Nature Reviews. Neurology 16 (11): 601–17.

Dijkgraaf, Feline E., Tiago R. Matos, Mark Hoogenboezem, Mireille Toebes, David W. Vredevoogd, Marjolijn Mertz, Bram van den Broek, et al. 2019. “Tissue Patrol by Resident Memory CD8+ T Cells in Human Skin.” Nature Immunology 20 (6): 756–64.

Fillatreau, Simon, Claire H. Sweenie, Mandy J. McGeachy, David Gray, and Stephen M. Anderton. 2002. “B Cells Regulate Autoimmunity by Provision of IL-10.” Nature Immunology 3 (10): 944–50.

Fitzpatrick, Zachary, Gordon Frazer, Ashley Ferro, Simon Clare, Nicolas Bouladoux, John Ferdinand, Zewen Kelvin Tuong, et al. 2020. “Gut-Educated IgA Plasma Cells Defend the Meningeal Venous Sinuses.” Nature 587 (7834): 472–76.

Frebel, Helge, Kirsten Richter, and Annette Oxenius. 2010. “How Chronic Viral Infections Impact on Antigen-Specific T-Cell Responses.” European Journal of Immunology 40 (3): 654–63.

Ghersi-Egea, Jean-François, Nathalie Strazielle, Martin Catala, Violeta Silva-Vargas, Fiona Doetsch, and Britta Engelhardt. 2018. “Molecular Anatomy and Functions of the Choroidal Blood-Cerebrospinal Fluid Barrier in Health and Disease.” Acta Neuropathologica 135 (3): 337–61.

Gray, Joshua I., and Donna L. Farber. 2022. “Tissue-Resident Immune Cells in Humans.” Annual Review of Immunology, January. https://doi.org/10.1146/annurev-immunol-093019-112809.

Häusser-Kinzel, Silke, and Martin S. Weber. 2019. “The Role of B Cells and Antibodies in Multiple Sclerosis,Neuromyelitis Optica, and Related Disorders.” Frontiers in Immunology 10 (February): 201.

Horns, Felix, Cornelia L. Dekker, and Stephen R. Quake. 2020. “Memory B Cell Activation, Broad Anti-Influenza Antibodies, and Bystander Activation Revealed by Single-Cell Transcriptomics.” Cell Reports 30 (3): 905–13.e6.

Jain, Rajiv W., and V. Wee Yong. 2021. “B Cells in Central Nervous System Disease: Diversity, Locations and Pathophysiology.” Nature Reviews. Immunology, December. https://doi.org/10.1038/s41577-021-00652-6.

Jarius, Sven, Friedemann Paul, Diego Franciotta, Patrick Waters, Frauke Zipp, Reinhard Hohlfeld, Angela Vincent, and >Brigitte Wildemann. 2008. “Mechanisms of Disease: Aquaporin-4 Antibodies in Neuromyelitis Optica.” Nature Clinical Practice. Neurology 4 (4): 202–14.

Jiang, Xiaodong, Rachael A. Clark, Luzheng Liu, Amy J. Wagers, Robert C. Fuhlbrigge, and Thomas S. Kupper. 2012. “Skin Infection Generates Non-Migratory Memory CD8+ TRM Cells Providing Global Skin Immunity.” Nature 483 (7388): 227–31.

Krumbholz, Markus, Tobias Derfuss, Reinhard Hohlfeld, and Edgar Meinl. 2012. “B Cells and Antibodies in Multiple Sclerosis Pathogenesis and Therapy.” Nature Reviews. Neurology 8 (11): 613–23.

Lanz, Tobias V., R. Camille Brewer, Peggy P. Ho, Jae-Seung Moon, Kevin M. Jude, Daniel Fernandez, Ricardo A. Fernandes, et al. 2022. “Clonally Expanded B Cells in Multiple Sclerosis Bind EBV EBNA1 and GlialCAM.” Nature, January. https://doi.org/10.1038/s41586-022-04432-7.

Lee, Dennis S.W., Olga L. Rojas, and Jennifer L. Gommerman. 2021. “B Cell Depletion Therapies in Autoimmune Disease: Advances and Mechanistic Insights.” Nature Reviews. Drug Discovery 20 (3): 179–99.

Li, Rui, Kristina R. Patterson, and Amit Bar-Or. 2018. “Reassessing B Cell Contributions in Multiple Sclerosis.” Nature Immunology 19 (7): 696–707.

MacLean, Andrew J., Niamh Richmond, Lada Koneva, Moustafa Attar, Cesar A. P. Medina, Emily E. Thornton, Ariane Cruz Gomes, et al. 2022. “Secondary Influenza Challenge Triggers Resident Memory B Cell Migration and Rapid Relocation to Boost Antibody Secretion at Infected Sites.” Immunity, March. https://doi.org/10.1016/j.immuni.2022.03.003.

Mazzitelli, Jose A., Leon C. D. Smyth, Kevin A. Cross, Taitea Dykstra, Jerry Sun, Siling Du, Tornike Mamuladze, Igor Smirnov, Justin Rustenhoven, and Jonathan Kipnis. 2022. “Cerebrospinal Fluid Regulates Skull Bone Marrow Niches via Direct Access through Dural Channels.” Nature Neuroscience, 1–6.

Meinl, Edgar, Markus Krumbholz, and Reinhard Hohlfeld. 2006. “B Lineage Cells in the Inflammatory Central Nervous System Environment: Migration, Maintenance, Local Antibody Production, and Therapeutic Modulation.” Annals of Neurology 59 (6): 880–92.

Neumeier, Daniel, Alessandro Pedrioli, Alessandro Genovese, Ioana Sandu, Roy Ehling, Kai-Lin Hong, Chrysa Papadopoulou, et al. 2021. “Profiling the Specificity of Clonally Expanded Plasma Cells during Chronic Viral Infection by Single-Cell Analysis.” European Journal of Immunology, November. https://doi.org/10.1002/eji.202149331.

Owens, Trevor, Ingo Bechmann, and Britta Engelhardt. 2008. “Perivascular Spaces and the Two Steps to Neuroinflammation.” Journal of Neuropathology and Experimental Neurology 67 (12): 1113–21.

Page, Nicolas, Sylvain Lemeille, Ilena Vincenti, Bogna Klimek, Alexandre Mariotte, Ingrid Wagner, Giovanni Di Liberto, Jonathan Kaye, and Doron Merkler. 2021. “Persistence of Self-Reactive CD8+ T Cells in the CNS Requires TOX-Dependent Chromatin Remodeling.” Nature Communications 12 (1): 1009.

Pollok, Karolin, Ronja Mothes, Carolin Ulbricht, Alina Liebheit, Jan David Gerken, Sylvia Uhlmann, Friedemann Paul, Raluca Niesner, Helena Radbruch, and Anja Erika Hauser. 2017. “The Chronically Inflamed Central Nervous System Provides Niches for Long-Lived Plasma Cells.” Acta Neuropathologica Communications 5 (1): 88.

Prinz, Marco, and Josef Priller. 2017. “The Role of Peripheral Immune Cells in the CNS in Steady State and Disease.” Nature Neuroscience 20 (2): 136–44.

Prüss, Harald. 2021. “Autoantibodies in Neurological Disease.” Nature Reviews. Immunology 21 (12): 798–813.

Qiu, Lin, Han Liu, Shuang Wang, Xiao-Hua Dai, Jian-Wei Shang, Xiao-Li Lian, Guan-Hua Wang, and Jun Zhang. 2021. “FKBP11 Promotes Cell Proliferation and Tumorigenesis via p53-Related Pathways in Oral Squamous Cell Carcinoma.” Biochemical and Biophysical Research Communications 559 (June): 183–90.

Ramesh, Akshaya, Ryan D. Schubert, Ariele L. Greenfield, Ravi Dandekar, Rita Loudermilk, Joseph J. Sabatino Jr, Matthew T. Koelzer, et al. 2020. “A Pathogenic and Clonally Expanded B Cell Transcriptome in Active Multiple Sclerosis.” Proceedings of the National Academy of Sciences of the United States of America 117 (37): 22932–43.

Sabatino, Joseph J., Jr, Anne-Katrin Pröbstel, and Scott S. Zamvil. 2019. “B Cells in Autoimmune and Neurodegenerative Central Nervous System Diseases.” Nature Reviews. Neuroscience 20 (12): 728–45.

Schenkel, Jason M., and David Masopust. 2014. “Tissue-Resident Memory T Cells.” Immunity 41 (6): 886–97.

Schwartz, Michal, Jonathan Kipnis, Serge Rivest, and Alexandre Prat. 2013. “How Do Immune Cells Support and Shape the Brain in Health, Disease, and Aging?” The Journal of Neuroscience: The Official Journal of the Society for Neuroscience 33 (45): 17587–96.

Shlesinger, Danielle, Kai-Lin Hong, Ghazal Shammas, Nicolas Page, Ioana Sandu, Andreas Agrafiotis, Victor Kreiner, et al. 2022. “Single-Cell Immune Repertoire Sequencing of B and T Cells in Murine Models of Infection and Autoimmunity.” bioRxiv. https://doi.org/10.1101/2022.02.07.479381.

Smets, I., and M. J. Titulaer. 2022. “Antibody Therapies in Autoimmune Encephalitis.” Neurotherapeutics: The Journal of the American Society for Experimental NeuroTherapeutics, January. https://doi.org/10.1007/s13311-021-01178-4.

Steinbach, Karin, Ilena Vincenti, Mario Kreutzfeldt, Nicolas Page, Andreas Muschaweckh, Ingrid Wagner, Ingo Drexler, Daniel Pinschewer, Thomas Korn, and Doron Merkler. 2016. “Brain-Resident Memory T Cells Represent an Autonomous Cytotoxic Barrier to Viral Infection.” The Journal of Experimental Medicine 213 (8): 1571–87.

Tan, Hyon-Xhi, Jennifer A. Juno, Robyn Esterbauer, Hannah G. Kelly, Kathleen M. Wragg, Penny Konstandopoulos, Sheilajen Alcantara, et al. 2022. “Lung-Resident Memory B Cells Established after Pulmonary Influenza Infection Display Distinct Transcriptional and Phenotypic Profiles.” Science Immunology 7 (67): eabf5314.

Tesfagiorgis, Yodit, Sarah L. Zhu, Rajiv Jain, and Steven M. Kerfoot. 2017. “Activated B Cells Participating in the Anti-Myelin Response Are Excluded from the Inflamed Central Nervous System in a Model of Autoimmunity That Allows for B Cell Recognition of Autoantigen.” The Journal of Immunology. https://doi.org/10.4049/jimmunol.1602042.

Urban, Stina L., Isaac J. Jensen, Qiang Shan, Lecia L. Pewe, Hai-Hui Xue, Vladimir P. Badovinac, and John T. Harty. 2020. “Peripherally Induced Brain Tissue-Resident Memory CD8+ T Cells Mediate Protection against CNS Infection.” Nature Immunology 21 (8): 938–49.

Vincenti, Ilena, Nicolas Page, Karin Steinbach, Alexander Yermanos, Sylvain Lemeille, Nicolas Nunez, Mario Kreutzfeldt,et al. 2022. “Tissue-Resident Memory CD8+ T Cells Cooperate with CD4+ T Cells to Drive Compartmentalized Immunopathology in the CNS.” Science Translational Medicine 14 (640): eabl6058.

Wang, Yan, Dianyu Chen, Di Xu, Chao Huang, Ruxiao Xing, Danyang He, and Heping Xu. 2021. “Early Developing B Cells Undergo Negative Selection by Central Nervous System-Specific Antigens in the Meninges.” Immunity 54 (12): 2784–94.e6.

Wilson, Emma H., Wolfgang Weninger, and Christopher A. Hunter. 2010. “Trafficking of Immune Cells in the Central Nervous System.” The Journal of Clinical Investigation 120 (5): 1368–79.

Yermanos, Alexander, Andreas Agrafiotis, Raphael Kuhn, Damiano Robbiani, Josephine Yates, Chrysa Papadopoulou, Jiami Han, et al. 2021. “Platypus: An Open-Access Software for Integrating Lymphocyte Single-Cell Immune Repertoires with Transcriptomes.” NAR Genomics and Bioinformatics 3 (2). https://doi.org/10.1093/nargab/lqab023.

Yermanos, Alexander, Daniel Neumeier, Ioana Sandu, Mariana Borsa, Ann Cathrin Waindok, Doron Merkler, Annette Oxenius, and Sai T. Reddy. 2021. “Single-Cell Immune Repertoire and Transcriptome Sequencing Reveals That Clonally Expanded and Transcriptionally Distinct Lymphocytes Populate the Aged Central Nervous System in Mice.” Proceedings of the Royal Society B: Biological Sciences 288 (1945): 20202793.

